# Cytoplasmic membrane thinning observed by interfacial dyes is likely a common effect of bactericidal antibiotics

**DOI:** 10.1101/2022.06.22.497132

**Authors:** Ashim Kumar Dubey, Taru Verma, Deepika Sardana, Balaram Khamari, Parvez Alam, Eswarappa Pradeep Bulagonda, Sobhan Sen, Dipankar Nandi

## Abstract

The lipid membrane is a fundamental part of life. However, the effects of different stresses on membranal integrity and physiology are less understood. Using novel 4-aminophthalimide-based membrane-specific dyes (4AP-Cn: n is carbon chain-length), aided with confocal microscopy, fluorescence spectroscopy, molecular dynamics simulations, and flow cytometry, we have studied stress-mediated changes in *E. coli* membranes. By exploiting the depth-dependent positioning and subsequent environmental sensitivity of the dyes, we have proposed a measure of antibiotic-induced membrane damage: the fluorescence Peak Maxima Difference (PMD) between 4AP-C9 and 4AP-C13. The ROS-influenced PMD quantifies cytoplasmic membrane thickness and measures sensitivity against most bactericidal antibiotics, depending upon the extent of lipid peroxidation. Importantly, we have verified this observation using antibiotic-sensitive and resistant clinical isolates of *E. coli* and ESKAPE pathogens like *K. pneumoniae* and *Enterobacter spp.* This study underscores the role of antibiotic-induced membrane changes; a critical issue given the increasing global antimicrobial resistance.

## Introduction

Lipid membranes are ubiquitous in all life. However, the inner workings of membranes, especially concerning stimuli responses, are not well understood (1). While a host of biophysical methods have helped understand aspects of membrane physiology, the scarcity of reliable fluorescence-based techniques (2) prompts the development of new live-cell lipid-specific fluorescent probes. Studying the properties of rigid gel-like phases of lipid/water interfaces, common in bacterial membranes, has been difficult (3). Here, we use novel interfacial polarity-sensitive dyes (4AP-Cn), which have 4-aminophthalimide attached to aliphatic chains (4), with *n* referring to the chain length. These dyes partition at different depths across the gel-phase lipid/water interface, depending on their aliphatic chain lengths and provide information on depth-dependent environmental polarity by their fluorescent peak shifts (4, 5). While the 4AP-Cn dyes have been characterised in DOPC/DPPC membranes (4, 5), their applications in living cells are yet to be studied. This study aims to understand bacterial membranes using this novel class of dyes.

Perturbations in the membrane affect bacteria significantly (6), making it crucial to understand their dynamics under various stresses. Antibiotic overuse, coupled with the slowdown in their development (7), has led to the rise of Antimicrobial Resistance (AMR). The global burden of bacterial AMR is over 1.2 million directly attributable deaths/year in 2019 (8), with estimates of up to 10 million AMR-related deaths/year by 2050 (9). The ability to accurately study membrane responses to antibiotic stress (10) enables significant diagnostic improvements. In this work, we have characterised the 4AP-Cn dyes mainly with the Gram-negative bacterium *E. coli.* We identified the dyes’ localisation to be primarily in the bacterial inner membrane via molecular dynamics simulations. Using different types of antibiotics in various bacterial species, we show a common effect of antibiotic action: cytoplasmic membrane thinning. This is likely caused by reactive species generation upon bactericidal antibiotic action leading to peroxidation of the cytoplasmic membrane lipids. We extend the use of these dyes to also study isolates of various clinically relevant bacteria and identified a robust, fluorescence-based methodology to detect antibiotic resistance profiles inexpensively and quickly.

## Results

### 4AP-Cn dyes stain bacterial membranes with high efficiency

To investigate whether 4AP-Cn dyes can stain bacterial membranes specifically and efficiently, we co-stained WT *E. coli* with the 4AP-Cn dyes and Nile Red, a fluorescent lipophilic dye that partitions into lipid membranes (11). Live WT *E. coli* cells were grown for 6 hrs (mid to late log phase) and stained with the dyes, followed by confocal imaging. The 4AP-Cn dyes used here have alkyl chains of 9 (4AP-C9) and 13 (4AP-C13) (Figures 1A and 1B). Significant colocalization was observed (Figures 1C and 1D), implying that the 4AP-Cn dyes are membrane dyes in live cells. Super-Resolution Radial Fluctuations (SRRF) analysis performed on 100 frames showed the same membrane colocalization (Figures 1C and 1D), implying that the membrane staining with the 4AP-Cn dyes was stable. There were two avenues of evidence to address the dyes’ membrane specificity – first, the SRRF analysis of stained bacteria over 500 frames (Figure 1F and 1G). The second was the separation of staining observed with the 4AP-Cn dyes and DRAQ, a far-red dye that stains only DNA (12). The co-staining of cells (Figure 1E) showed that the two dyes were separated, implying the membrane specificity of the 4AP-Cn dyes.

**Figure 1.**
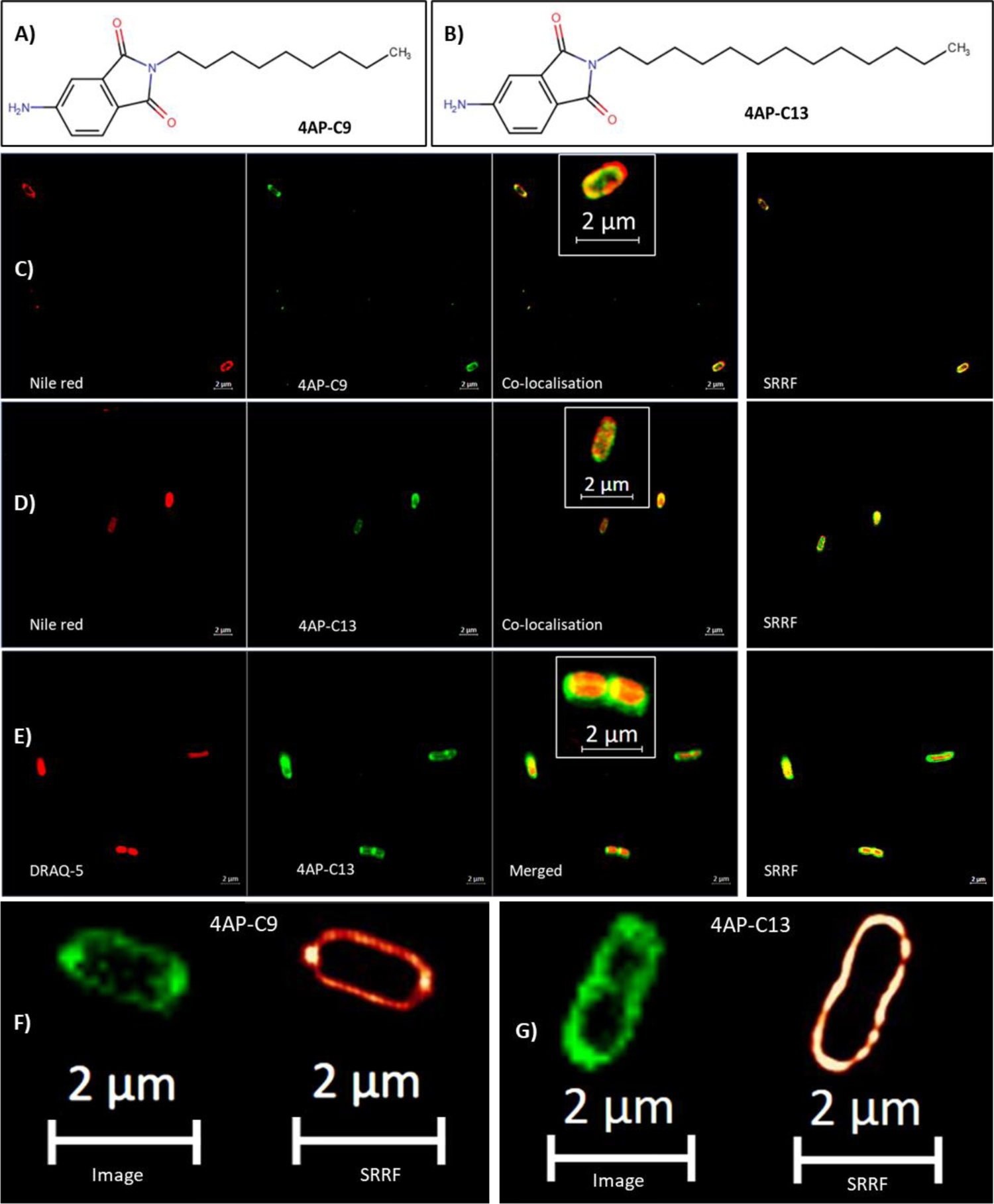
The interfacial dyes 4AP-C9 and 4AP-C13 stain bacterial membranes. The 4AP-Cn dyes consist of the fluorescent 4-aminophthalimide (4AP) attached with an aliphatic chain. The dyes used in this study have aliphatic chains of length 9 – 4AP-C9 (A) and length 13 – 4AP-C13 (B). Confocal microscopy (under AiryScan mode) was used to show the co-localisation of the lipid membrane dye Nile Red with 4AP-C9 (C) and 4AP-C13 (D) in live Wild Type *E. coli* K12 MG1655 cells grown for 6 hours. The separation of staining between the membrane dye 4AP-C13 and the DNA staining dye, DRAQ is shown (E). In (C), (D) and (E), column 1 and column 2 show the dyes as separate channels, while column 3 shows both channels together, along with a zoomed single-cell inset image for clarity. Column 4 shows the merged false colour image from SRRF analysis of 100 frames. The specificity of the 4AP-Cn dyes is abundantly clear by their localisation in the membrane, as shown in the comparison of a single frame image and the SRRF analysis of drift corrected 500 frame data set, for staining with 4AP-C9 (F) and 4AP-C13 (G).

Next, we compared the staining of live cells, permeabilised cells, and PFA fixed cells (13). Permeabilization was performed using Polymyxin B Nonapeptide (PMBN), a derivative of Polymyxin B (PMB) deficient in antibacterial activity, which still can permeabilize the outer membrane (14). Fixation by PFA completely abrogates cellular functions (13) and permeabilises the membranes (15). Upon confocal imaging of live and fixed cells, we did not observe any significant difference in their morphology. However, the fraction of stained cells was higher after fixation. Using flow cytometry, we observed that the 4AP-Cn dyes stained live cells better than Nile red (Supplementary Figure 1). Permeabilization by PMBN increased staining by 4AP-C13 but did not affect staining by the others, suggesting that mild permeabilization was not enough. However, in all dyes, fixation significantly increased the percentage of stained cells (Supplementary Figure 1), suggesting that the total membrane permeabilization and disrupted efflux allow for greater dye uptake (16). Importantly, these results confirm that the 4AP-Cn dyes are membrane specific, with enhanced staining upon fixation.

### 4AP-Cn dyes are interfacial in bacterial membranes with their fluorescence depending on temperature-dependent membrane fluidity

We compared 4AP-Cn staining with bacterial cells’ staining with Laurdan, a fluorescence probe that shows emission changes based on membrane fluidity (17). We compared the fluorescence changes in the 4AP-Cn dyes on modulating the membrane fluidity via rapid cooling of bacteria grown for 3 hrs (early to mid-log) and 6 hrs (mid to late-log). We observed that 4AP-C9 was unaffected by temperature changes. In contrast, 4AP-C13 was significantly blue-shifted, potentially due to the temperature-induced fluidity reduction, across time points (Figures 2A - C). Changes in membrane fluidity were also verified using Laurdan fluorescence (Figure 2D) and quantified via GP measurements (Equation 1), which showed an inverse correlation with fluidity. The temperature-induced reduction in membrane fluidity was as expected for cells grown for 3 and 6 hrs (Figures 2E and 2F). The spectral changes observed here suggest that 4AP-Cn intercalation in bacterial cells depends upon membrane fluidity, with a gel-like phase allowing for a stable, deeper intercalation. This gives a quantifiable measure for membrane properties: the Peak Maxima Difference (PMD) between the fluorescence of 4AP-C9 and 4AP-C13 at low temperatures.

**Figure 2.**
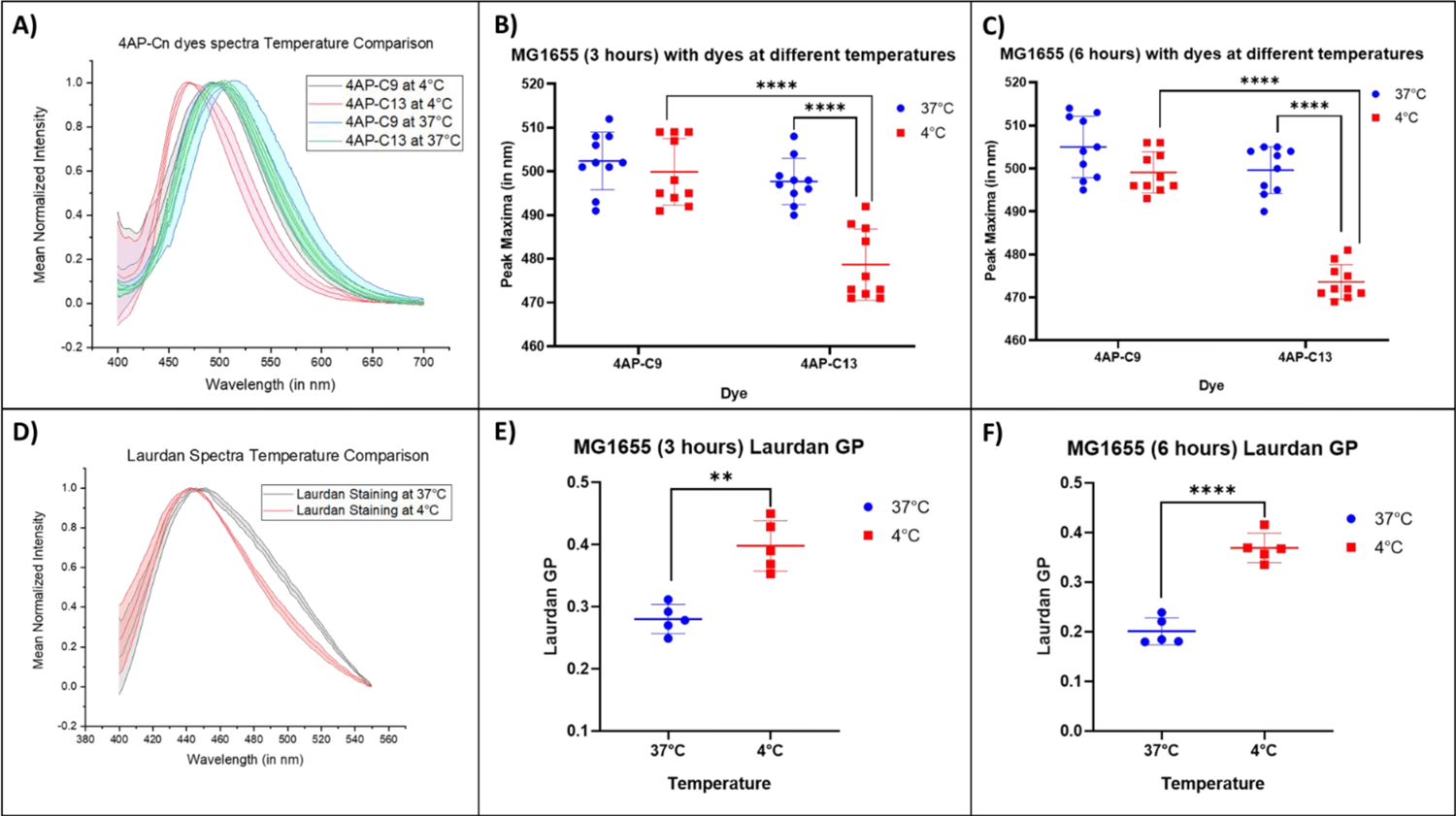
4AP-Cn dyes are interfacial, and their fluorescence is dependent upon temperature –. The first row shows (A) the 6-hour representative spectra comparisons, followed by the peak maxima comparison between the 4AP-Cn dyes across temperatures, for untreated live Wild Type *E. coli* MG1655 bacteria grown for (B) 3 hours and (C) 6 hours. The second-row shows (D) the 6-hour representative spectra comparisons for Laurdan, followed by the Laurdan GP comparison in live bacteria across temperatures for (E) 3 hours and (F) 6 hours. In (B) and (C), statistical analysis has been performed using two-way ANOVA, while in (E) and (F), it was performed using Welch’s t-test, where * indicates *P* < .05; **, *P* < .01; ***, *P* < .001 and ****, *P* < .0001.

### The 4AP-Cn dyes interact primarily at the lipid/water interfaces of the bacterial inner membrane

To distinguish whether insertion of 4AP-C9 and 4AP-C13 occurs in the bacterial inner membrane (IM) or outer membrane (OM), MD simulations were performed in PL:PL IM and LPS:PL OM with and without the dyes (Supplementary Table 1 and Supplementary Figure 2). The dynamics of the dyes’ interaction with the IM and OM were studied at 30°C using all-atom MD simulations to observe the location (depth) distributions, hydration state and angle distributions of dyes at the lipid/water interface (Figure 3). The ordering of the lipid chains is unaffected by the incorporation of the dyes, as seen by the SCD of various lipid chains in the absence and presence of the 4AP-Cn dyes (Supplementary Figure 3).

**Figure 3.**
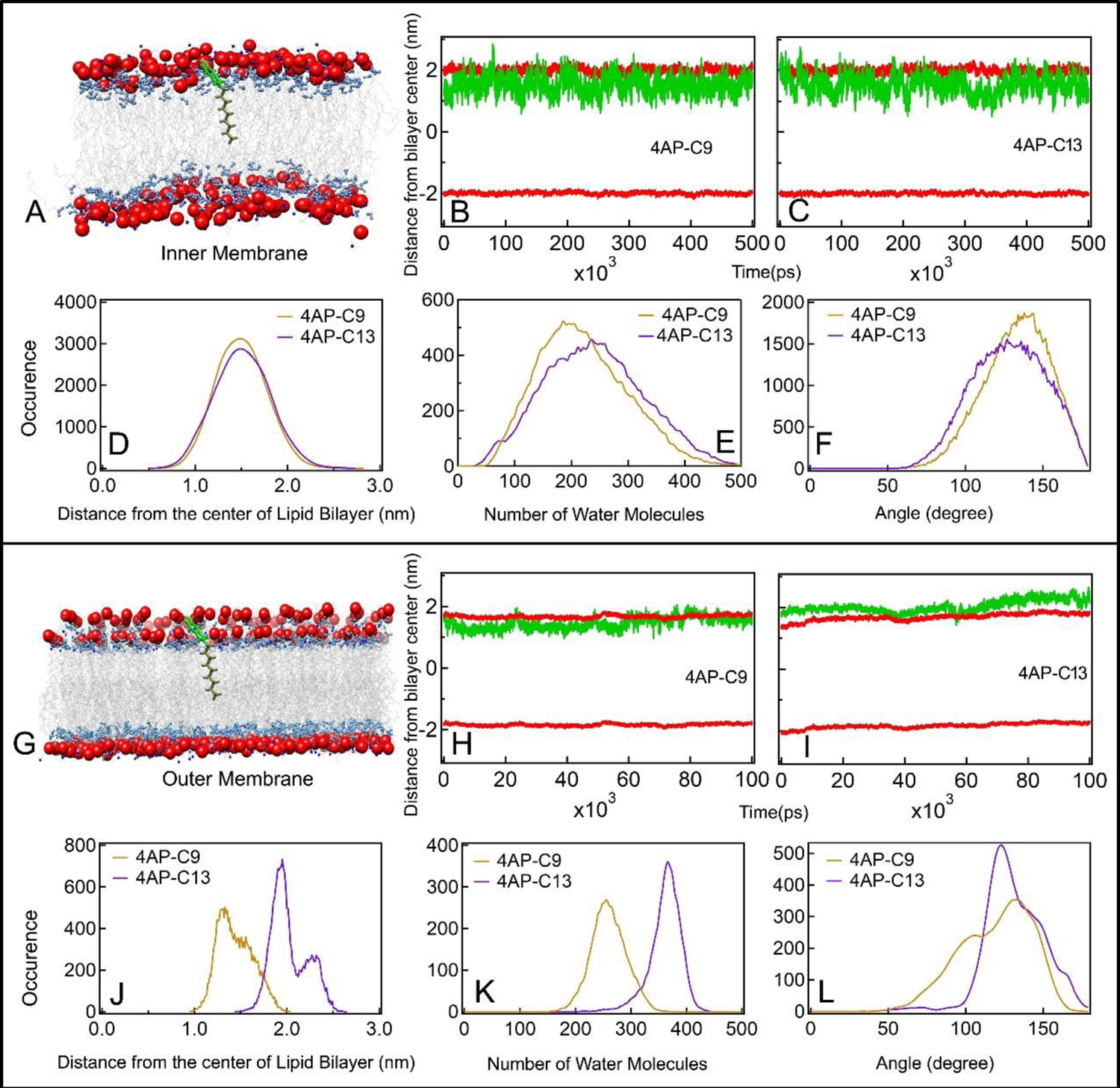
The interfacial properties of the 4AP-Cn dyes arise from the staining of the cytoplasmic inner membrane. Representative snapshots of the MD simulation show the most probable position of the 4AP-Cn dye (4AP-C9 shown for representation here) inside the (A) symmetric PL:PL Inner Membrane (IM) and (G) asymmetric LPS:PL Outer Membrane (OM) at 303.15 K (Water molecules in both IM and OM, and the core region and O Antigen in OM have not been shown for clarity. Full snapshots are shown in Supplementary Figure 2). Fluctuations of positions of centre-of-mass of 4AP moiety in 4AP-C9 and 4AP-C13 throughout 500 ns in IM (B and C) and 100 ns in OM (H and I) are plotted, respectively. Distributions of z-position fluctuations of the dyes in the IM (D) and OM (J), their hydration states in the IM (E) and OM (K), and angle distributions in the IM (F) and OM (L) are also plotted. The colour code used is as follows: blue – nitrogen atoms; red – phosphorous atoms, light blue – ester groups, grey – alkyl tails of lipids; green – 4AP moiety; tan – aliphatic chain of 4AP-Cn dyes.

The snapshot of the IM/4AP-Cn system and the fluctuations of z-positions of 4AP-C9 and 4AP-C13 were visualized (Figures 3A-C). Distributions of position fluctuations of the two dyes showed that the dyes’ average positions are similar in the IM (Figure 3D), possibly due to high fluidity, like the experimental observations at 37 °C. Fluctuations in the dyes’ distributions in the OM, however, showed that the dyes were shifted, with 4AP-C13 significantly exposed toward the water phase when inserted in the LPS leaflet of OM (Figure 3).

Distributions of the number of water molecules within 1 nm of 4AP-moiety within which the dipolar water govern the local solvation (polarity) of the dyes (4) (Figure 3E), and orientation of the dyes (Figure 3F) were very similar in the IM, however, in the OM, 4AP-C13 was highly exposed to water (Figure 3K), with the dye orientations sharp and shifted (Figure 3L). Reconciling the experimental and simulation data, the 4AP-Cn dyes are mainly adsorbed within the IM of bacterial cells, as the LPS leaflet is not thick enough to allow for the deeper intercalation of the 4AP-C13 dye necessary for the spectral changes observed in the experiments.

### 4AP-C13 with DRAQ5 can be used to detect morphological changes in bacteria treated with antibiotics

Next, we studied the dyes’ ability to detect morphological changes due to antibiotic treatment, by using three antibiotics, PMB, tetracycline and ciprofloxacin, based on their different modes of action (Supplementary Table 2). In all three antibiotics, WT *E. coli* cells were grown for 6 hrs at subinhibitory concentrations (0.7XMIC) of the antibiotics (18), stained, and imaged.

We observed significant morphology changes across antibiotic treatments compared to untreated bacterial cells (Figure 4A): cell rounding in PMB treated cells (Figure 4B), cell lengthening and increased curvature in tetracycline treated cells (Figure 4C) and filamentation and disruption of DNA localisation in ciprofloxacin treated cells (Figure 4D). SRRF analysis on longer time-series data (500 frames) showed that the untreated bacterial cells did not show any specific structural defect in the membranes (Figure 4D), whereas the PMB treated cells were rounded (Figure 4E). There were noticeable membrane invaginations under tetracycline treatment (Figure 4F) and significant membranal disruptions and intercalations into the filamented cell under ciprofloxacin treatment (Figure 4G). Bright field images also showed similar membrane changes (Supplementary Figure 4), showing the observed phenotypes to be common across stained and unstained cells.

**Figure 4.**
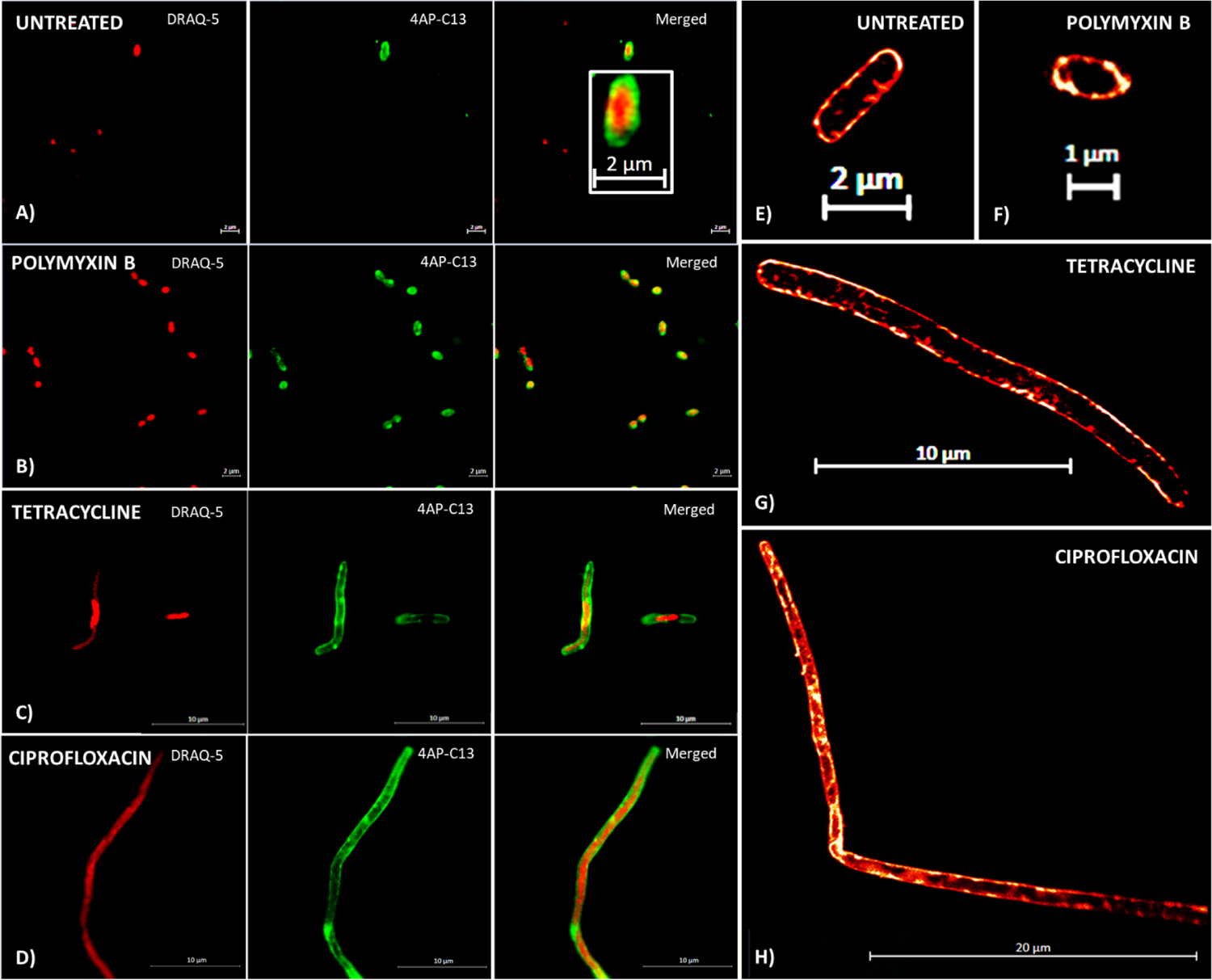
Morphological impact of antibiotics on bacteria can be clearly visualized using the 4AP-C13 and DRAQ5 dye system –. (A, B, C, D) Confocal microscopy under AiryScan mode is used to show the morphological changes in Wild Type *E. coli* MG1655 on growth for 6 hours in the presence and absence of antibiotics, using the membrane dye, 4AP-C13 and the DNA staining dye, DRAQ5. (E, F, G, H) The membrane morphology changes are shown with greater clarity by the performance of SRRF on 500 frames of drift corrected time series confocal images on cells stained with 4AP-C13. Visualized here, are the live Wild Type *E. coli* cells, which are (A, E) untreated, or treated with 0.7XMIC of (B, F) Polymyxin B, (C, G) Tetracycline and (D, H) Ciprofloxacin. In (A), (B), (C) and (D), columns 1 and 2 show the two dyes, DRAQ5 and 4AP-C13, as separate channels, while column 3 shows both channels together.

### Differing membrane effects of antibiotics are quantifiable by the fluorescence PMD of the 4AP-Cn dyes

Fluorescence PMDs between the two dyes were studied to understand the modulation of membrane properties in bacteria treated with different antibiotics. Cells were grown for 3 hrs and 6 hrs under treatment with ciprofloxacin (Figure 5A), tetracycline (Figure 5B) and PMB (Figure 5C), fixed, stored, stained and their spectra measured. The spectra after fixation and overnight storage were indistinguishable from those from rapidly cooled live cells (data not shown). The data was fitted with a negative Hill function, considering that the antibiotic-induced PMD reduction can be regarded as “inhibition” of the PMD (19).

**Figure 5.**
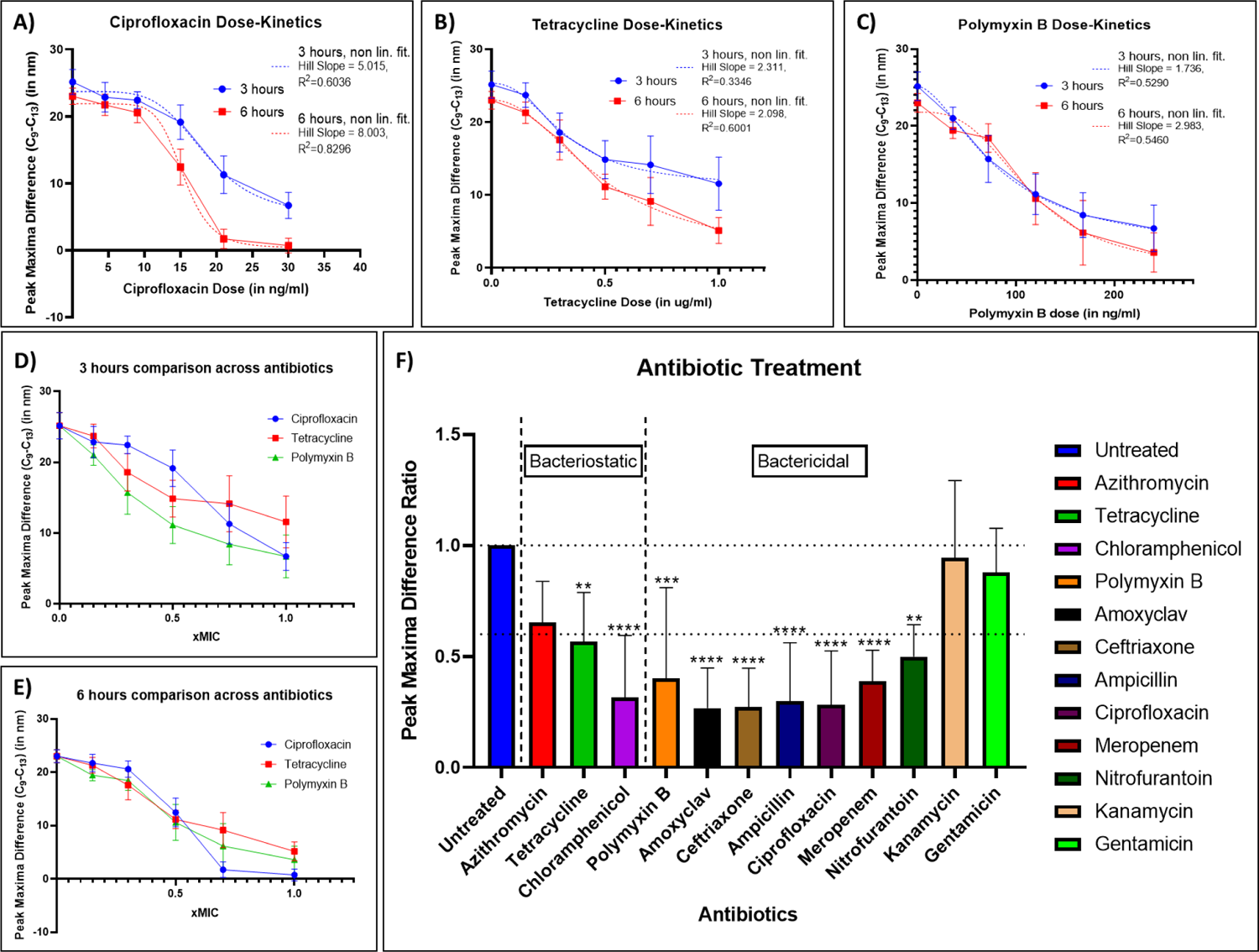
*E. coli* under different antibiotic treatments show demonstrable and separable effects on the peak maxima difference (PMD) of the 4AP-Cn dyes. Wild Type *E. coli* MG1655 was grown for 3 and 6 hours in the presence of different concentrations of various antibiotics, and the bacteria were then fixed and stored overnight. The cells were then stained with the 4AP-Cn dyes, and the peak maxima differences (PMDs) were observed. The first row shows the dose-dependent curve of the peak maxima difference of the 4AP-Cn dyes across time, in presence of the antibiotics – A) Ciprofloxacin, B) Tetracycline and C) Polymyxin B. The same data is then represented as a comparison across antibiotics, separated by the time points – D) 3 hours and E) 6 hours. Hill Function curves have been fit to the data in A), B), and C), to understand incremental antibiotic effects on the membrane, with the parameters of fitting mentioned alongside. F) This figure compares the peak maxima difference ratio (PMDR) for cells grown for 6 hours in the presence of ∼0.7xMIC of antibiotics, both bacteriostatic and bactericidal. The bacteriostatic antibiotics used here are azithromycin, tetracycline, and chloramphenicol, while the bactericidal antibiotics used are polymyxin B, amoxyclav (amoxicillin-clavulanic acid), ceftriaxone, ampicillin, ciprofloxacin, meropenem, nitrofurantoin, and the aminoglycosides – kanamycin and gentamicin. In F) statistical analysis was performed here using two-way ANOVA, where * indicates *P* < .05; **, *P* < .01; ***, *P* < .001 and ****, *P* < .0001. The data shown in each panel is representative of 7 replicates.

Under ciprofloxacin treatment, the PMD curve had a high Hill slope at 3 and 6 hrs. At 3 hrs, the PMD was reduced, the Hill’s slope was lower, and the fit was poorer. (Figure 5A) The high Hill’s coefficient suggested that ciprofloxacin affected the membrane indirectly by downstream effects, like the induction of reactive species (20–23). There has been (some contradictory) evidence of ciprofloxacin action on membranes (24–26), including cardiolipin rearrangement (27); however, the results here verify ciprofloxacin’s membrane effects. Under tetracycline treatment, the PMD curve had a lower Hill’s slope for both time points, implying a more direct membrane action (Figure 5B). The peak maxima reduction was less than that under ciprofloxacin treatment, implying a lower membrane disruption, in a slow process with heterogeneity reducing over time. This delayed, direct membrane effect of tetracycline was unexpected (28). However, tetracycline affecting membranes directly in a mechanism independent of ribosome inhibition has been recently characterised (29). Under PMB treatment, the PMD curve had a much lower Hill’s slope (Figure 5C), indicating the antibiotic’s direct mechanism of action on bacterial membranes (30, 31). However, the fit was not good for both the time points, with the curves being similar, which suggested that the membrane effects may be rapid (32).

The same data was grouped by time of growth rather than the antibiotic used. The analysis of growth for 3 hrs (Figure 5D) and 6 hrs (Figure 5E) revealed that the PMDs and the variations in the antibiotics were best observed at the 6-hour time point for the ∼0.7XMIC of that antibiotic-strain combination. Consequently, this particular combination has been selected for subsequent studies.

### PMD reduction of the 4AP-Cn dyes arises due to cytoplasmic membrane thinning

The 4AP-Cn dyes’ PMD arises because of differences in depth-dependent polarity of 4AP-C9 and 4AP-C13 in membranes (4, 5). The disappearance of the PMD is likely to arise from two possibilities: high membrane fluidity and membrane thinning. To understand which of the two explanations is more likely, comparisons were performed with Laurdan which showed significant temperature-induced fluidity decrease across time points. At the 3-hour point, there was no significant difference across antibiotic treatments at both temperatures (Supplementary Figure 5A). At the 6-hour point, there was a significant fluidity decrease at 37°C by the antibiotics, with ciprofloxacin and PMB showing large drops. At 4°C, there was a significant fluidity difference between the untreated bacteria and the ciprofloxacin and PMB treated bacteria (Supplementary Figure 5B). This data suggests no significant fluidity increase occurred on antibiotic treatment in our experimental conditions. Instead, there was a reduction in membrane fluidity, especially with ciprofloxacin. This suggests that membrane thinning is crucial in the antibiotic-induced effect observed by the dyes.

### Antibiotic action may be causing cytoplasmic membrane thinning in bacteria as a common mechanism

To study whether PMD reduction is a common mechanism of antibiotic action, we performed the fluorescence spectroscopy assays on different bacteria-antibiotic combinations under the same testing conditions (Supplementary Table 2). The concentrations of each antibiotic used (Supplementary Table 3) were selected by broth-microdilution experiments to keep the OD_600_∼0.2 at the measurement time point of 6 hours (Supplementary Figure 10A) to normalize for growth related effects. Most antibiotic treatments showed significant peak maxima difference ratio (PMDR: Equation 2) reductions (Figure 5F), except for Azithromycin and the aminoglycosides. Grouping the antibiotics by their growth impact on bacteria showed some degree of PMDR reduction by most bacteriostatic antibiotics, though there is significant variation. On the other hand, bactericidal antibiotics show significant PMDR reduction uniformly, (except aminoglycosides) suggesting a common bactericidal antibiotic action on membranes.

### Membrane thinning upon antibiotic treatment is independent of antibiotic-induced cell size changes

Next, we tested whether antibiotic-induced changes in cell size or granularity can explain the observations across the antibiotic treatment by performing flow cytometry on different antibiotic-treated bacteria (33). We compared the forward scatter (cell size) and side scatter (granularity) (34) (SSC-A vs FSC-A curves) for all the different antibiotics used in our study (Supplementary Figure 6). We observed that the changes induced by the antibiotics were apparent, with the curve shifts upon treatments with azithromycin, tetracycline, the β-lactams, or the cell size reduction with PMB being especially notable. We compared the median FSC-A across treatments to measure cell size (Supplementary Figure 6N), with significant differences under antibiotic treatment; however, the cell size and PMD reduction data had insignificant correlation.

### Reactive species generation and membrane lipid peroxidation leads to membrane thinning by bactericidal antibiotics

There is significant evidence for reactive species generation being a general mechanism of bacterial death under the action of bactericidal antibiotics (20, 35, 36). We investigated the roles of ROS by using the cell-permeable dye CellRox ® Orange. The bacterial metabolism regulators like NADH dehydrogenase regulate ROS generation (37), whose positioning explains the colocalization of the ROS and membrane dyes in our untreated sample (Supplementary Figure 7A). This spatial proximity allows for increased ROS amounts potentially causing cytoplasmic lipids peroxidation. Confocal imagery showed that this proximity remains upon antibiotic treatment (Supplementary Figures 7B-D).

We compared the histograms of the stained and unstained cells across the antibiotic treatments and observed minimal fluorescence changes upon staining, for untreated cells (Figure 6A) or tetracycline treated cells (Figure 6B), with a significant increase upon ciprofloxacin treatment (Figure 6C). To quantify the ROS induction across the different antibiotic treatments, we used a Median Fluorescence Index (MFI) based methodology (Supplementary Figure 7E). Quantifying cellular ROS revealed that bacteriostatic antibiotics did not induce ROS (20, 35, 36). Bactericidal antibiotics significantly increased the cellular ROS, except for the aminoglycosides and PMB (Figure 6D). The order of ROS amounts for bactericidal antibiotics was gentamicin ∼kanamycin< PMB< nitrofurantoin< ceftriaxone< ciprofloxacin< ampicillin< meropenem< amoxyclav, with clear shifts observed in the histograms (Figure 6E). The PMDR and the ROS amounts ratio showed a high correlation, especially for bactericidal antibiotics (Figure 6F).

**Figure 6.**
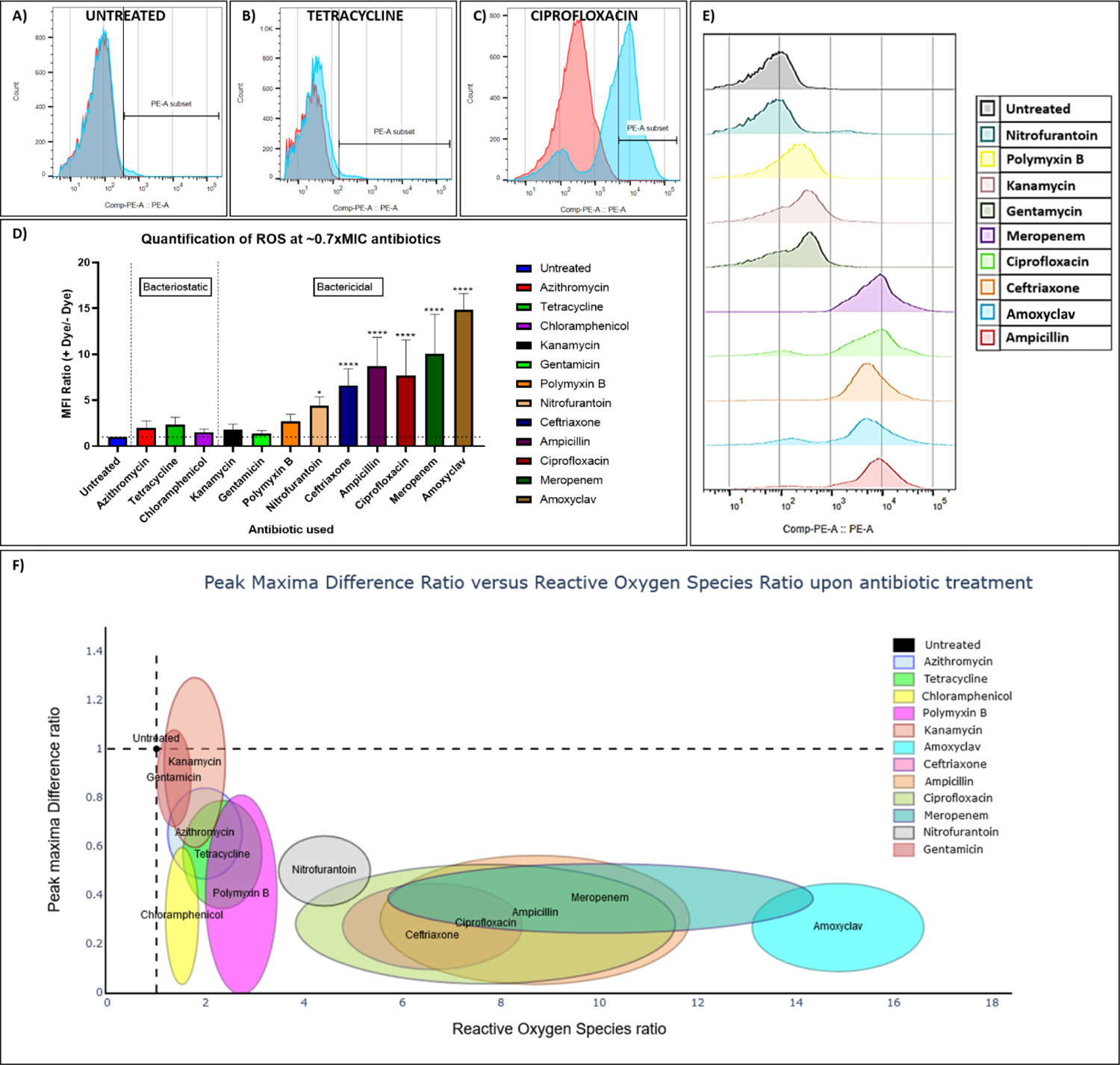
The peak maxima difference ratio (PMDR) reduction on antibiotic action is a function of ROS induction by bactericidal antibiotics. A representative histogram for the unstained and stained cells with the ROS dye CellROX Orange using flow cytometry, for A) Untreated cells, B) Tetracycline treated cells, and C) Ciprofloxacin treated cells. D) This figure compares the quantification of the ROS MFI Ratio, for cells grown for 6 hours in the presence of ∼0.7xMIC of antibiotics. The bacteriostatic antibiotics used here are azithromycin, tetracycline, and chloramphenicol, while the bactericidal antibiotics used are polymyxin B, amoxyclav (amoxicillin-clavulanic acid), ceftriaxone, ampicillin, ciprofloxacin, meropenem, nitrofurantoin and the aminoglycosides, kanamycin, and gentamicin. E) This figure shows a representative overlay of the flow cytometry histograms as obtained from the bactericidal antibiotic-treated cells stained with CellROX Orange. F) This figure compares the peak maxima difference ratio with the ROS ratio, across the different antibiotic treatments, via ellipses for each antibiotic with the Mean +/- SD as the bounds for each measure. The data in each quantification panel is representative of 7 replicates. Statistical analysis was performed here using one-way ANOVA, where * indicates *P* < .05; **, *P* < .01; ***, *P* < .001 and ****, *P* < .0001.

The results from the ROS quantification suggested the possibility of lipid peroxidation. Therefore, we performed flow cytometry with the lipid peroxidation dye, BODIPY-C11 (38, 39), which can determine the extent of peroxidation in a membrane by fluorescing differently upon peroxidation. We tested for peroxidation with antibiotics representative of each group: tetracycline-bacteriostatic which did not induce ROS; gentamicin-bactericidal which did not induce ROS and ciprofloxacin-bactericidal which significantly induced ROS. The histograms show a negligible increase in the median fluorescence of the cells in the presence of the dye for untreated cells (Figure 7A), tetracycline treated cells (Figure 7B), or gentamicin treated cells (Figure 7C) with a significant increase upon ciprofloxacin treatment (Figure 7D). This was further quantified using a Median Fluorescence Index (MFI) based ratio-metric methodology (Supplementary Figure 8), showing similar results (Figure 7E). Notably, small amounts of peroxidised lipids can significantly affect the IM (40). This suggests that lipid peroxidation-induced cytoplasmic membrane thinning may be common in bactericidal antibiotic action, which can be detected using the 4AP-Cn dyes.

**Figure 7.**
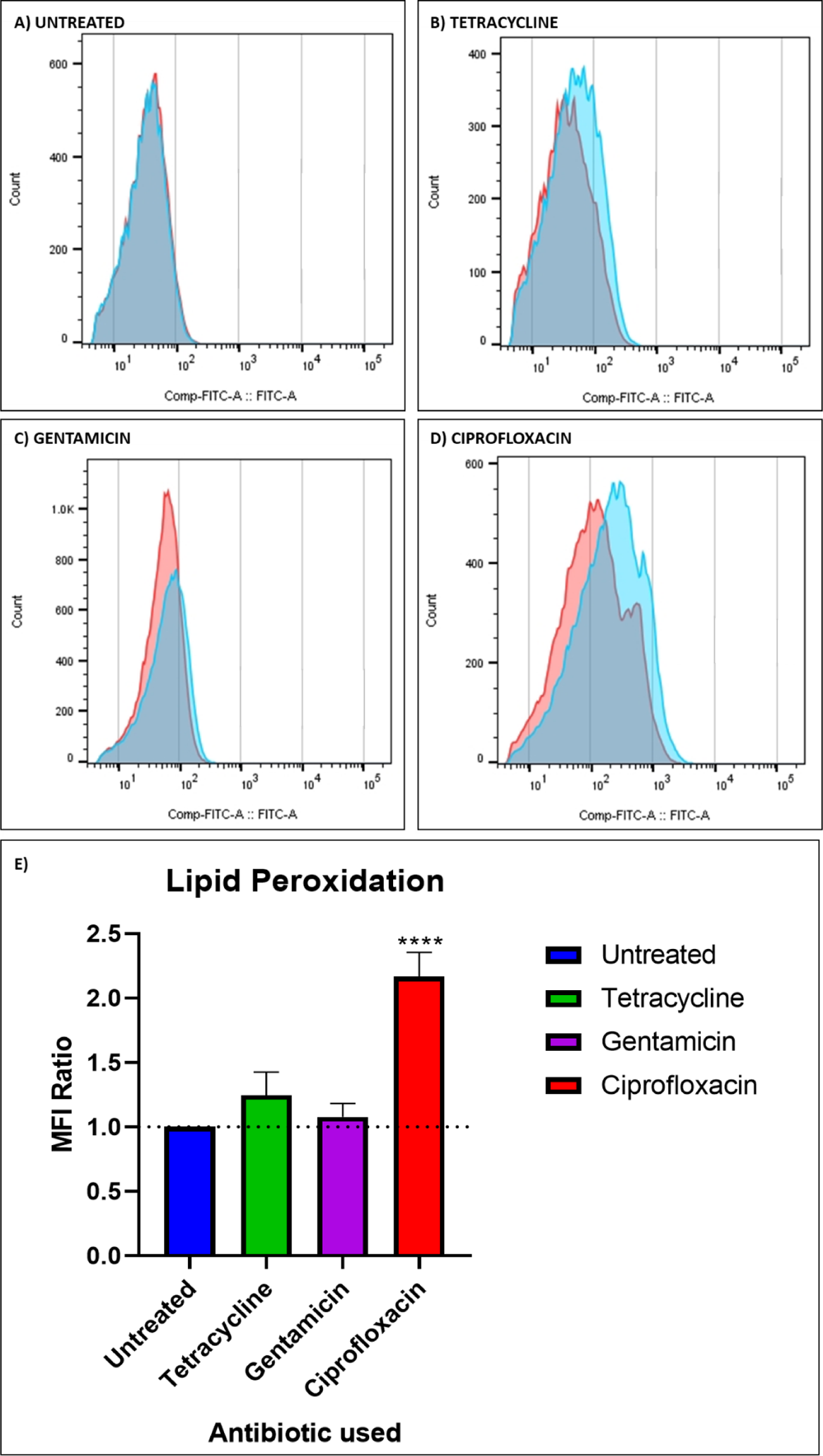
Lipid Peroxidation due to higher ROS induction may be responsible for membrane thinning, leading to the antibiotic-induced peak maxima difference reduction. A representative histogram for the unstained and stained cells with the lipid peroxidation dye BODIPY-C11 as obtained using flow cytometry is shown, for A) Untreated cells, B) Tetracycline treated cells, C) Gentamicin treated cells, and D) Ciprofloxacin treated cells. E) This figure compares the quantification of lipid peroxidation, via the MFI Ratio, for cells grown for 6 hours in the presence of ∼0.7xMIC of the antibiotics tetracycline, gentamicin, and ciprofloxacin. The data shown here is representative of 7 replicates. Statistical analysis was performed here using one-way ANOVA, where * indicates *P* < .05; **, *P* < .01; ***, *P* < .001 and ****, *P* < .0001.

### Antioxidants mitigate the PMDR reduction upon antibiotic action

To validate our prior findings, we studied the dyes’ fluorescence in bacteria grown in the simultaneous presence of ciprofloxacin and antioxidants. For this, we used glutathione, which is a known scavenger of reactive metabolite by-products (20). While an averaged spectra in the absence of glutathione for untreated and ciprofloxacin treated cells showed differences between the spectra (Figure 8A), supplementation by 1mM glutathione (Figure 8B) and 5mM glutathione (Figure 8C), showed a dose-dependent rescue. Although glutathione treatment did not cause any change, there was a significant, dose-dependent rescue of the PMDR for cells grown in the presence of glutathione and ciprofloxacin (Figure 8D). We also used the antioxidant N-Acetyl Cysteine (NAC) (43) to confirm that this was not a glutathione-specific process. Although there was no NAC-dependent peak maxima change, a significant, dose-dependent rescue of the PMDR upon growth under ciprofloxacin treatment in the presence of NAC was observed (Figure 8E). Finally, to monitor the effects of attenuation of lipid peroxidation, we used α-tocopherol, which mitigates lipid peroxidation in eukaryotes (41). The antioxidant did not affect the PMDR by itself. Low antioxidant concentrations led to a lesser rescue, whereas a high concentration led to a significant rescue of the PMDR reduction (Figure 8F).

**Figure 8.**
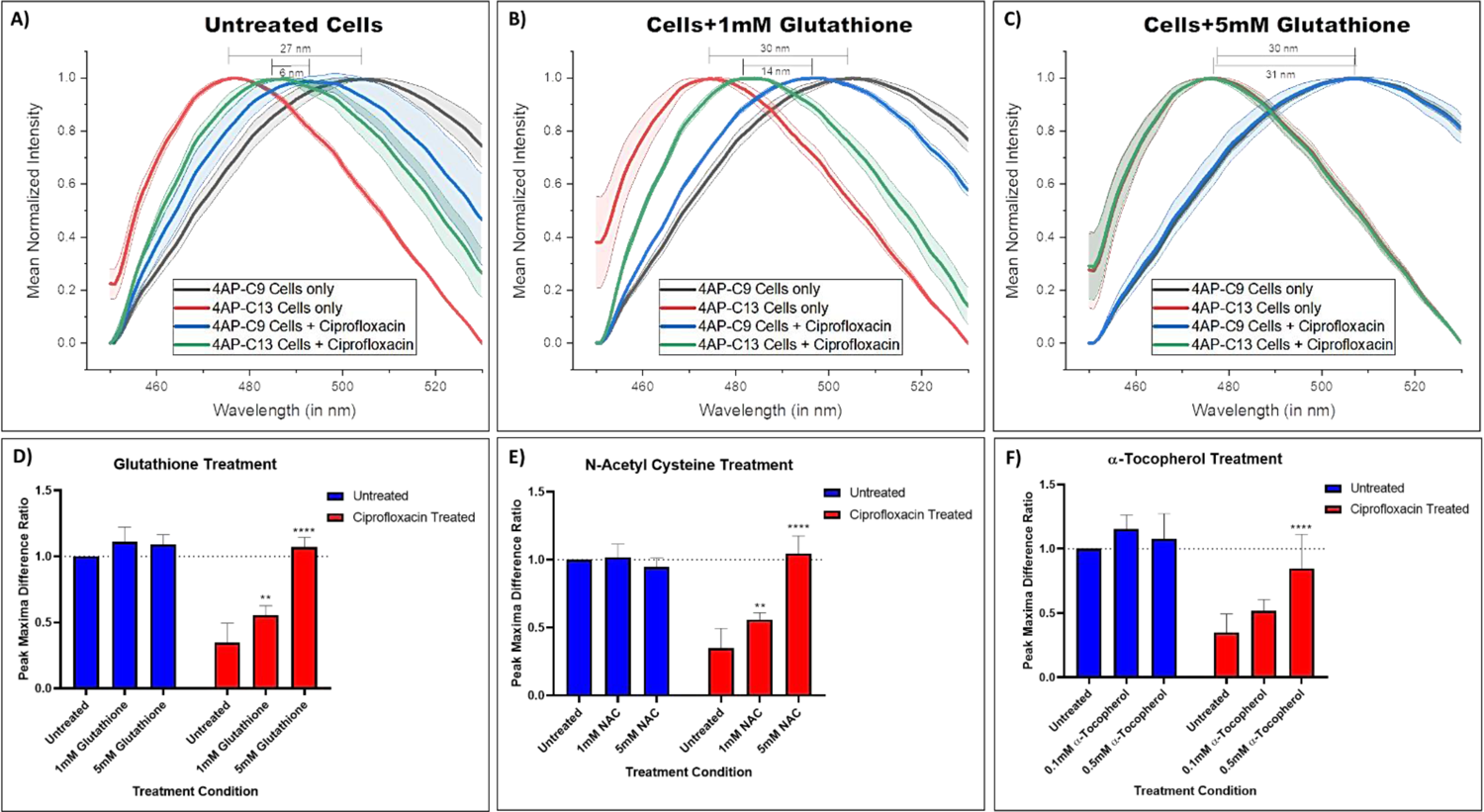
Quenching of Reactive Oxygen Species by different ROS Quenchers leads to a rescue of the observed peak maxima difference ratio reduction (PMDR), in a dose-dependent manner. The fluorescent spectra of 4AP-C9 and 4AP-C13 treated cells in the presence and absence of the antibiotic ciprofloxacin, along with the peak maxima difference between the dyes, has been shown for differing concentrations of the ROS quencher Glutathione – A) untreated cells (control), B) 1mM Glutathione and C) 5mM Glutathione, showing a dose-dependent rescue in PMDR. The quantification of the dose-dependent rescue in the presence of ROS quenchers of the PMDR reduction on ciprofloxacin treatment is visualized for D) Glutathione, E) N-Acetyl Cysteine and F) α-Tocopherol. The data shown in each panel is representative of 7 replicates. Statistical analysis was performed here using two-way ANOVA, where * indicates *P* < .05; **, *P* < .01; ***, *P* < .001 and ****, *P* < .0001.

The antioxidants also rescued growth under ciprofloxacin treatment, seen by the dose-dependent changes in the OD_600_ values (Supplementary Figure 10B). Together, this data suggests that the quenching or scavenging of ROS, either inside the cell or at the membrane, leads to a significant rescue of the bacterial growth and PMDR reduction, suggesting that cytoplasmic membrane thinning under bactericidal antibiotic treatment is influenced by the ROS generation, and the ensuing membrane lipid peroxidation.

### Cytoplasmic membrane thinning is a measure of bacterial sensitivity to antibiotic action

It was important to address whether membrane thinning is an accurate measure of sensitivity to antibiotics and whether it could be used to determine the resistance profile of a bacterium. Therefore, we performed our fluorescence assay across mechanisms of antibiotic resistance. It has been found that *lon* defective mutants of *E. coli* have excess accumulation of the MarA protein, which upregulates the AcrAB-TolC efflux pump and makes them inherently resistant to a host of antibiotics (42–45). We observed that the 4AP-Cn dyes’ PMDR was significantly less for WT bacteria than the Δ*lon* strain under ciprofloxacin and tetracycline treatment, unlike PMB treatment (Supplementary Figure 9A). Thus, the membrane effects observed upon antibiotic treatment depend on susceptibility as genotypically resistant bacteria don’t show the PMDR reduction. Furthermore, phenotypic mechanisms of antibiotic resistance were tested using Sodium salicylate (NaSal). NaSal is known to bind the MarR protein, leading to the de-repression of MarA expression (46). NaSal is also known to induce copper ion-dependent oxidation of MarR cysteine residues (47). Cells were grown in the presence of 3mM NaSal alongside ciprofloxacin to study the effects of phenotypic resistance. Interestingly, we saw a ratio reduction on just NaSal, which may have arisen from ROS induction (48). We saw a significant rescue in the PMDR in the presence of NaSal and ciprofloxacin (Supplementary Figure 9B), showing that phenotypic resistance mechanisms show the same differentiable impact on antibiotic action.

A similar experiment was also performed with *Staphylococcus aureus*, a Gram-positive bacterium. Vancomycin sensitive and resistant strains of *S. aureus* were grown in the presence and absence of ciprofloxacin and vancomycin for 6 hrs. While the growth studies matched our expectations, as seen by the OD_600_ of the different treatment conditions (Supplementary Figure 9D), vancomycin treatment did not significantly reduce the PMDR for the sensitive isolate. However, upon ciprofloxacin treatment, we observed a significant PMDR reduction in both strains (Supplementary Figure 9C).

Together, these studies demonstrate the potential of this combination of dyes to function as a potent measure of resistance for bacterial samples. While the resistant samples show no significant PMDR reduction, the sensitive samples do, implying that cytoplasmic membrane thinning is a function of bacterial sensitivity.

### Clinical isolates of pathogens show cytoplasmic membrane thinning in an antibiotic sensitivity dependent manner

It was essential to verify whether the phenotypes observed in the *E. coli* lab strain could be confirmed in clinical isolates across antibiotics and bacterial species. The antibiotic dose and time point to match our criteria were found by a 96-well plate checkerboard assay with clinical bacterial isolates, with the measurement of the OD_600_ of the bacteria across time points. Using this, time points, isolates and antibiotic concentrations were determined, with attempts to ensure two sensitive and two resistant isolates per antibiotic-bacteria combination. The samples were then grown in a 50ml culture in a 250ml flask with appropriate antibiotic concentrations (Supplementary Table 4), fixed with 4% PFA after growth, stained, and analysed.

Initially, clinical isolates of *E. coli* were studied, and their PMDRs were quantified. Resistant and sensitive isolates were designated with a colour-coded method by their OD_600_ readings – >∼50% growth reduction in the presence of the antibiotic compared to its absence is designated sensitive (Supplementary Figure 11A-F). The gentamicin treated cells did not show a PMDR reduction. While all the samples obtained were sensitive to nitrofurantoin and PMB, there was no drastic decrease in the PMDR. The samples treated with the bactericidal antibiotics that induce high ROS, i.e., ciprofloxacin, meropenem and amoxyclav showed a noticeable PMDR reduction in sensitive cells and no difference in resistant cells (Figure 9A).

**Figure 9.**
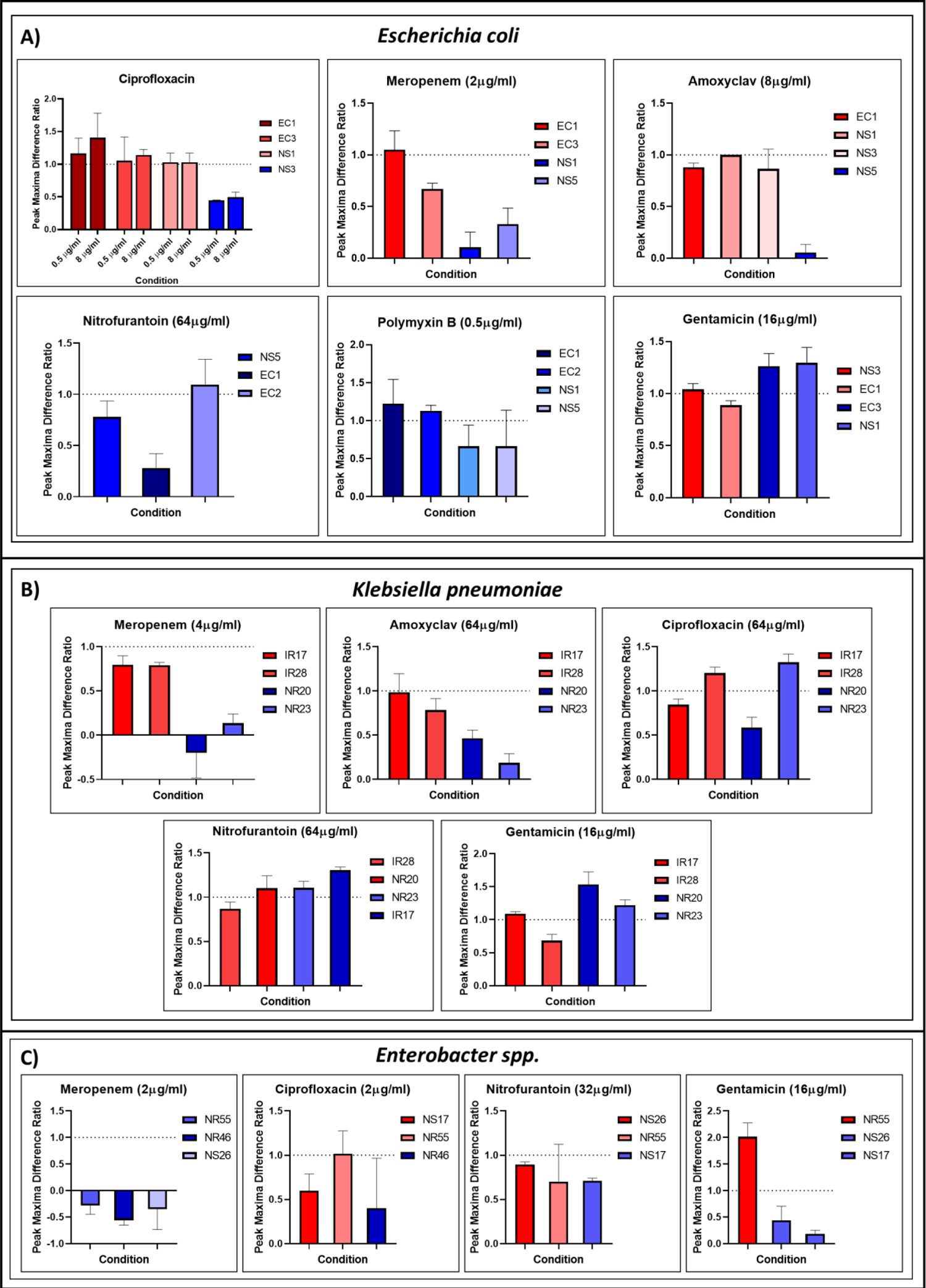
Clinical isolates of *Escherichia coli, Klebsiella pneumoniae* and *Enterobacter spp.* show membrane thinning for various antibiotic treatments. The peak maxima difference ratio (PMDR) of clinical isolates of *Escherichia coli* (A), *Klebsiella pneumoniae* (B) and *Enterobacter spp.* (C), of differing antibiotic sensitivities, were measured after they were grown in the presence of relevant concentrations of selected antibiotics. The isolates marked red are resistant to the antibiotic, while those marked blue are sensitive. Sensitivities and resistances are designated based on OD_600_ reductions of the isolate on antibiotic treatment – >∼50% reduction in the presence of the antibiotic compared to its absence is designated sensitive. Each data point is representative of 2 technical replicates.

We also investigated whether the dyes can identify sensitivities and resistances in other bacteria of clinical importance, like some ESKAPE pathogens (49). *Klebsiella pneumoniae*, *Enterobacter* spp. and *Acinetobacter baumannii* clinical strains were obtained after fixation with similar conditions of a few resistant and sensitive isolates, based upon the OD_600_ values (Supplementary Figure 12B-D). Among the strains, *Acinetobacter baumannii* isolates did not show the expected results across antibiotics (Supplementary Figure 12A). The clinical isolates of *K. pneumoniae* showed activity significantly similar to that of the *E. coli* samples. Treatment with nitrofurantoin and gentamicin did not show a PMDR reduction in sensitive cells, and ciprofloxacin treatment had just one sensitive isolate showing an effect. Treatment with meropenem and amoxyclav showed noticeable PMDR reduction in sensitive cells, and not so in resistant cells (Figure 9B). With *Enterobacter* spp., while nitrofurantoin treated samples did not show a sensitivity based peak maxima reduction; however, treatment with meropenem and ciprofloxacin showed the expected phenotype, with gentamicin unexpectedly showing a PMDR reduction (Figure 9C). Notably, despite the different antibiotics used, the PMDR reduction of the 4AP-Cn dyes could separate resistant and sensitive isolates of even some clinically relevant pathogens.

## Discussion

In this study, we characterised a novel class of membrane dyes, the 4AP-Cn dyes, and identified experimental conditions that lead to the efficient utilisation of their depth-dependent positioning to study membrane changes in bacteria. We have characterised a novel, fluidity-independent, fluorescence-based measure of antibiotic-induced membrane damage. This damage arises from reactive species generated upon bactericidal antibiotic treatment interacting with the IM lipids and peroxidising them, causing membrane damage and thinning. This can then be detected by interfacial dyes of different intercalations, like 4AP-C9 and 4AP-C13, with the observations being unlikely to be strictly growth related: all antibiotics reduce growth, but the cytoplasmic membrane thinning is observed mainly in the bactericidal antibiotics (Figure 5). We have created a mechanistic illustration for membrane changes in a gram-negative bacterium in the presence and absence of antibiotics (Figure 10).

**Figure 10.**
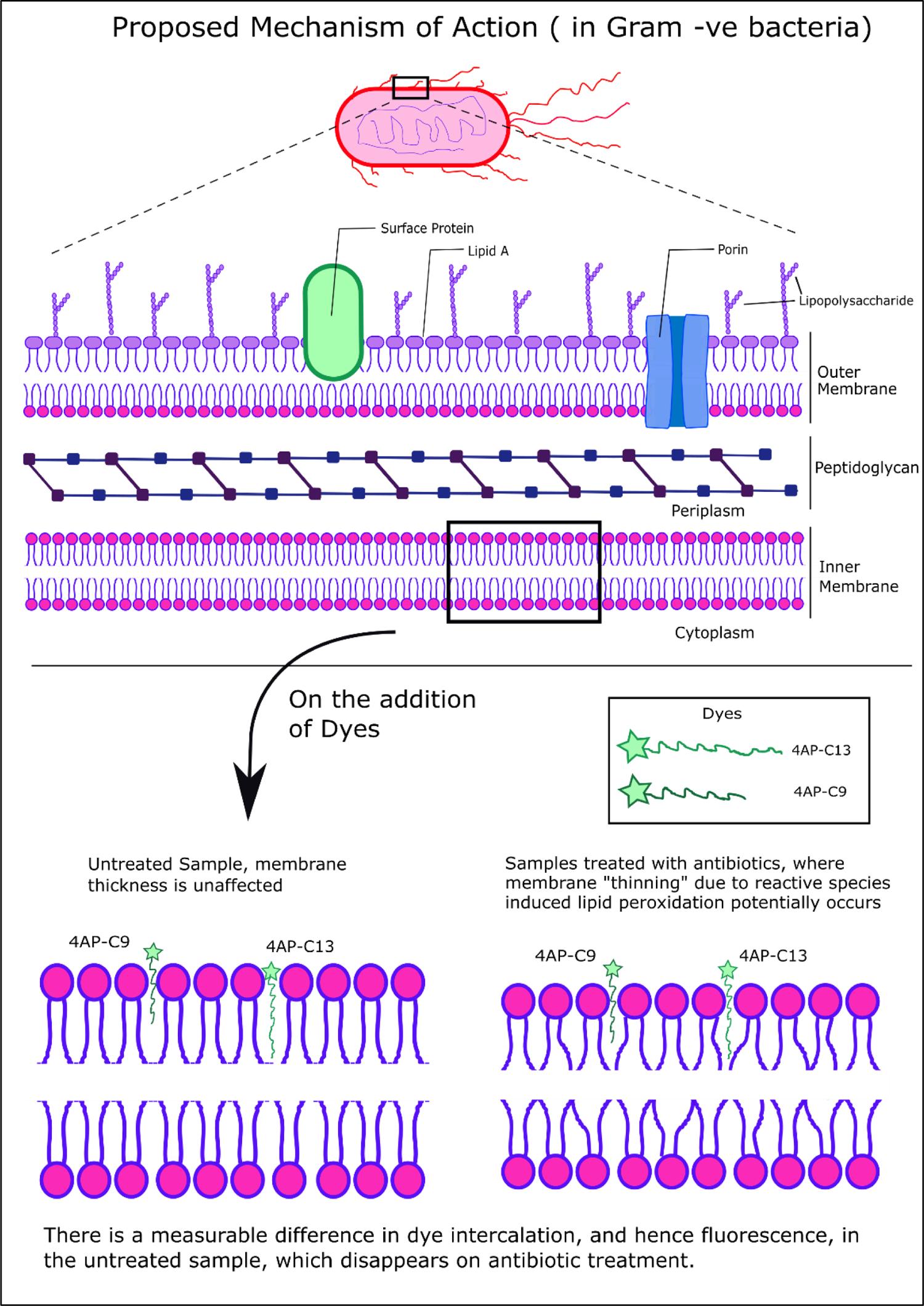
Proposed model to explain the mechanism of action that leads to the observation of a reduction in the peak maxima difference ratio (PMDR) of the fluorescence of the 4AP-Cn dyes under bactericidal antibiotic treatment. This figure first shows a general Gram-negative bacterium, followed by a zoom into the protective double membrane structure. The subsequent section shows the proposed mechanism of action – intercalation of the 4AP-Cn dyes most likely, primarily occurs in the inner membrane of Gram-negative bacteria. In the untreated sample, the dyes intercalate to slightly different positions, owing to the differing chain lengths. The difference in the positioning of the 4AP head group leads to a measurable fluorescence PMDR, whereas, in the antibiotic-treated sample, the reactive species generation leads to the peroxidation of the inner membrane lipids, leading to membrane disruption, hence “thinning”, leading to the 4AP-C13 dye being “pushed” out, thereby causing the loss of the fluorescence PMDR observed before.

The interfacial positioning nature of the 4AP-Cn dyes makes them very attractive to help understand membrane properties, given the deeper intercalation into gel-like membranes for increased carbon chain lengths. This effect, however, is not very significant in a fluid phase lipid–water interface (4). This implies that high fluidity can lead to a negligible PMD between 4AP-C9 and 4AP-C13, as was seen at 37°C in *E. coli*, due to 4AP-C13’s labile positioning (Figure 2). Furthermore, 4AP-C13 has an aliphatic chain 13 carbons long, similar to dyes like Dil-C12, which are used to selectively intercalate into the fluid pars of the membrane (50). To understand fluidity changes, we have used Laurdan, which primarily intercalates into the IM, though OM staining cannot be ruled out (51). Staining being cytoplasmic, however, matches our results (Supplementary Figure 5) – under PMB treatment, IM rigidity was increased, as seen in MD simulations that showed PMB’s ability to stiffen the IM (52). There have been few studies on antibiotic-induced membrane fluidity changes under ciprofloxacin (53) and tetracycline treatments (40). However, to the best of our knowledge, not much work has been done in the experimental conditions used here.

Apart from the possibility of heightened fluidity, the other possibility for diminished PMD upon treatment is “membrane thinning”, which has two possible origins: pressure induced thinning or via other biophysical interactions; and biochemical interactions that damage the membrane, making the nonpolar protective layer, i.e., the membrane, thinner. Upon thinning, 4AP-C13 is pushed out, as the IM leaflet has a thickness similar to the length of 4AP-C13. The dyes are unable to intercalate into the inner leaflet of IM. This has been characterised earlier, with 4AP-C12 in model DOPC/DPPC membranes (4). Bacteria can adapt to temperature changes over time (54), with significant effects on sustained membrane fluidity changes (55). However, rapid cooling down has been shown to significantly reduce membrane fluidity (56) without other adaptations. This makes our experimental protocol ideal for understanding the stress-mediated membrane thickness changes, as the low temperature removes the impact of fluidity. Our data demonstrate that cytoplasmic membrane thinning plays a significant role in antibiotic stress, as detected by the 4AP-Cn dyes (Figure 5).

Reactive species generation has been characterised as a general mechanism of death due to the action of bactericidal antibiotics (20, 35, 36). ROS and lipid peroxidation induction were quantified via fold change comparisons of stained and unstained cells in the presence and absence of antibiotics to handle antibiotic-induced autofluorescence changes (57). The ability of ciprofloxacin to induce ROS in bacteria has been shown before (20–23). High induction of ROS in cells treated with β-lactams has been explained recently, showing lethality under β-lactam action to be influenced by a rapid increase in anabolic and catabolic processes, leading to a large-scale accumulation of reactive by-products (58). The low amounts of ROS upon aminoglycoside action at the time studied explains the lack of a PMDR reduction. While ROS has been hypothesised to be necessary for aminoglycoside lethality (59), recent studies have shown that the antibiotic uptake modulated by the PMF dominates over ROS (60). The extent of PMDR reduction matches closely with ROS induction during treatment with bactericidal antibiotics (Figure 6). The significant increase in lipid peroxidation upon treatment with subinhibitory concentrations of ciprofloxacin is relevant as it provides a mechanistic aspect to our results (Figure 7). Lipid peroxidation has been shown to cause membrane thinning in MD simulations and experiments across different lipid bilayers (61) by the tendency of the oxidised lipids to bend towards the water interface. There are membrane disruptions at low concentrations of peroxided lipids, with increasing peroxidation causing more adverse effects. The antioxidants rescue the PMDR reduction (Figure 8), confirming the importance of ROS in the observed phenotype. PMDR reductions in bacteriostatic antibiotic treatments, however, also suggest for ROS-independent mechanisms of membrane thinning (Figure 5F).

There is significant evidence that *lon* mutants of *E. coli* have enhanced antibiotic resistance (42–44). The utilisation of the Δ*lon* strain and NaSal (62) gave us a direct correlation between antibiotic sensitivity and PMDR reduction. Also, clinical isolates of different bacterial species treated with antibiotics showed similar phenotypes for most combinations (except for *A. baumannii* isolates). While the β-lactams and fluoroquinolones showed the expected phenotypes across isolates, others like nitrofurantoin did not. This may be due to some antibiotics inducing lower amounts of ROS which do not lead to membrane damage that is significantly detectable by the 4AP-Cn dyes. Most of the variations observed here can be explained as clinical isolates differ from lab strains, with variable cell membrane thickness and antibiotic responses. Since we hypothesise that cytoplasmic membrane thinning is causing the peak maxima reduction, it stands to reason that there would exist a sweet spot of membrane thickness of the untreated cells where the dye combination is most likely to show the effect of treatments. This corresponds to the deeper intercalating dye (4AP-C13) having a length comparable to the lipids of the membrane (the phospholipids of the IM, having a length of ∼18 carbons), so that the leaflet is about as thick as the length of the intercalating dye, allowing membrane thickness changes to be reflected. If the lengths/thickness are not comparable, then the following will occur: when the membrane is too thin, the intercalating dye would already be in the pushed-out state, leaving a negligible or negative PMD, leading to the inability to observe membrane thinning, as seen under MD with LPS (Figure 3). Similarly, a thicker membrane would show a PMD under no treatment, but the dyes would be unlikely to pick up any membrane thinning because the intercalating dye would not be pushed out, leading to no observable change. This could be tackled by using dyes of lower aliphatic lengths in the former case and longer lengths in the latter, making the fluorescence spectroscopy of the 4AP-Cn dyes a finely tuneable method to study membrane properties near the lipid water interface.

Since cytoplasmic membrane thinning in sensitive bacteria depends on bactericidal antibiotic damage, one can design diagnostics of antibiotic resistance with quick turnovers using only the 4AP-Cn dyes’ fluorescence to help separate antibiotic resistance and sensitive infections with high specificity. Outputs in the lab were achieved rapidly after getting samples (∼24 samples in 2 hours). This is relevant as the urgency imposed by the rise of AMR necessitates gauging resistance in bacterial infections quickly, not just for the patient but also for antibiotic stewardship (63). Finally, the successful utilisation of a family of tuneable interfacial dyes to study the impact of stresses on cellular membranes is a novel concept. It opens the field of membranal studies to newer frontiers, potentially helping understand or even resolve some biomedical challenges, like AMR.

## Materials and Methods

### Bacterial Strains

*E. coli* WT MG1655 was obtained from the Coli Genetic Stock Centre, Yale University. The Δ*lon* strain in a chloramphenicol background was made and characterised in previous studies. (42, 43) The *S. aureus* strains were obtained from the lab of Prof. Jayanta Chatterjee, Molecular Biophysics Unit, IISc. At the same time, the clinical isolates of *E. coli*, *K. pneumoniae*, *Enterobacter spp.* and *A. baumannii* were fixed and generously provided to us by Dr B.E. Pradeep of Sri Sathya Sai Institute of Higher Learning, Puttaparthi, Andhra Pradesh.

### Antibiotic MIC Calculation

MIC of antibiotics with WT *E. coli* MG1655 was calculated using E-tests and spotting assays for ciprofloxacin (32 ng/ml), tetracycline (1050 ng/ml) and PMB (250 ng/ml), and the values obtained are close to those in literature. (43, 64, 65) For this study, the highest concentration taken for each antibiotic was: ciprofloxacin (30 ng/ml), tetracycline (1000 ng/ml) and PMB (240ng/ml). Broth macro dilution was used to supplement the data obtained from spotting assays and calculate the MICs for the other antibiotics and bacterial species.

### Sample Preparation

From a plate of bacteria, a primary inoculum was made from a single colony, grown overnight in Luria-Bertani (LB) medium consisting of 10 g of tryptone (Himedia Laboratories, Mumbai, India), 10 g of NaCl (Merck, Mumbai, India) and 5 g of yeast extract (Himedia) per litre at 37°C with constant shaking at 160 rpm, with requisite antibiotics for mutants (42, 43). The secondary culture was made with the following conditions – 10 ml LB in a 50 ml falcon, 0.2% OD2 normalised primary culture + antibiotics as required, at 37°C, 160 rpm. Appropriate amount of culture was centrifuged at 5000g for 10 minutes and resuspended in 1 mL of 1xPBS by gentle pipetting. We either proceeded for fixation or dye staining based on the condition being tested.

### Fixation

For fixation, an appropriate number of cells (∼10^7^ CFU) were resuspended in 1.5ml 4% PFA solution (13) and kept on ice for 30 to 45 mins. This incubation allowed fixation even at low-temperature conditions. This solution was then pelleted down, and the cells were then washed with 1x PBS to remove any residual PFA. The fixed cells were then resuspended in 1ml PBS for further processing and kept at 4°C until their usage.

### Materials for 4AP-Cn Synthesis

4-Aminophthalamide (4-AP) and alkyl bromides (nonyl bromide and tridecyl bromide) were purchased (TCI Chemicals, Chennai, India). 4-AP was recrystallised before using it to synthesise 4-AP probes (4AP-Cn) from an ethanol/water mixture. Anhydrous potassium carbonate (K_2_CO_3_) and silica gel were obtained (Thermo Fisher Scientific, Waltham, Massachusetts, USA). Water (HPLC grade) and ethanol (HPLC) were obtained from Merck. Ethyl acetate and hexane were obtained from Rankem. Acetonitrile (UV-spectroscopy grade) was obtained from Spectrochem, Mumbai, India.

### Synthesis of 4AP-Cn probes

4-AP-Cn were synthesised using a one-step reaction of 4-Aminophthalimide with alkyl bromide (nonyl bromide and tridecyl bromide, respectively), as discussed previously (4, 5). Nuclear magnetic resonance was used to characterise the 4-AP-Cn molecules (4AP-C9 and 4AP-C13). The NMR data are as follows:

**4AP-C9** ^1^HNMR (500MHz, CDCl_3_, 298K):0.8(t,3H);1.2(m,12H); 1.6(t,2H); 3.6(t,2H); 6.8(dd,1H);7.0(d,1H);7.5(d,1H);4.4(s,2H).

**4AP-C13** ^1^HNMR (500MHz, CDCl_3_, 298K): 0.9(t,3H);1.2(m,20H); 1.6(t,2H); 3.6(t,2H); 4.3(s, 2H); 6.8(dd,1H);7.0(d,1H);7.6(d,1H)

### Simulation Methods

#### Starting structures and parameters of 4AP-C*n*

The initial coordinates for 4AP-C9 and 4AP-C13 probes were obtained from energy optimized structures of the probes using Gaussian 09 software (66) at Hartree-Fock (HF) level using 6-31G^∗^ basis set (4). To calculate excited state charges of probes, the electrostatic potential (ESP) was generated on an optimized 4AP-C*n* structure at CIS/6-31G^∗^ level using Gaussian 09. Two-stage restrained electrostatic potential (RESP) charge methodology (67) was utilized to calculate the excited state atomic partial charges of 4AP-C*n.* The first charge optimization is done by applying a hyperbolic restraint of 0.0005 on non-hydrogen atoms, keeping the rest of the atoms free. In the second stage, charges are re-optimized and refit using hyperbolic restraint of 0.001 on non-hydrogen atoms, which freeze charges on all atoms except for methylene (CH2) and methyl group (CH3) of the carbon chains of 4AP-C*n* probes. General Amber Force Field (GAFF) (68) parameters were used to generate force-field parameters and topologies of 4AP-C*n* probes using in antechamber module of AMBER-18 (69), which were then converted into GROMACS format using python based ACPYPE (AnteChamber Python Parser Interface) script (70).

#### Starting structures and parameterization of lipids

Starting coordinates for all the membrane systems were generated from Membrane Builder (71, 72) available in CHARMM-GUI (73) using CHARMM36 lipid force field (74–77). The OM of gram-negative bacteria is an asymmetric bilayer where the outer leaflet mainly consists of LPS (lipopolysaccharides), and the inner leaflet contains different types of phospholipids (PLs) (77, 78). The inner membrane (IM) is symmetric and contains phospholipids of similar composition in both the leaflets. In our study, the inner leaflet of OM and both leaflets of IM consisted of a mixture of 1-palmitoyl(16:0)-2-palmitoleoyl(16:1 cis-9)-phosphatidylethanolamine (PPPE), 1-palmitoyl(16:0)-2-vacenoyl(18:1 cis-11)-phosphatidylglycerol (PVPG), and 1,10-palmitoyl-2,20-vacenoyl cardiolipin (PVCL2) in a ratio of 15:4:1 (PPPE : PVPG : PVCL2) (Supplementary Figure 2). The outer leaflet of OM is composed of the rough LPS with lipid A and R1 core with O-antigen polysaccharide. The lipid A of *E. coli* LPS consists of six amide/ester-linked fatty acids with a length of either 14 or 12 carbons (77, 78), with the details of the system used in this study summarised in Supplementary Table 1. 150 mM KCl was added to maintain the physiological ionic conditions. For the LPS system, Ca^2+^ ions were added to neutralise LPS-A lipids. TIP3P water model was used for all the simulations (79).

### MD Protocol

All lipid systems were first minimized and equilibrated with gradually reducing positional and dihedral restraints in six steps using the protocol recommended by the membrane builder in CHARMM-GUI. Then the 4AP-C*n* dyes were incorporated at the lipid/water interface of these equilibrated bilayers such that carbon chains of the dyes are oriented along the alkyl tails of lipids inside the leaflet. The lipid bilayers along the dyes were again energy minimized and equilibrated in NVT followed by NPT. The production simulations were performed at 303.15 K (30.15 °C) in NPT ensemble. During equilibration, semi-isotropic pressure coupling was used for lipid bilayers and LINCS algorithm to restraint bonds containing hydrogen (80). A constant atmospheric pressure of 1 bar was maintained using Parrinello-Rahman barostat with isothermal compressibility of 4.5 × 10^-5^ bar^-1^ and coupling constant of 5 ps, and Nose-Hoover thermostat (81). Particle Mesh Ewald was used to treat the long-range electrostatic interactions (82). MD simulation trajectories were saved at every 5 ps. Trajectories were analysed using analysis modules of GROMACS 2019. VMD (83) and Chimera (84) were used for the analysis and visualisation. The details of simulated systems and their total production runs are provided in Supplementary Table 1. The angle of inclination of the 4AP-moiety of the dyes with respect to the bilayer normal was calculated by drawing a vector connecting N2 and C2 atoms of 4AP-moiety and normal to lipid bilayer.

The effect of incorporation of dyes on the lipid order was investigated by calculating the carbon-deuterium order parameter, SCD, of the lipid chains in absence and presence of dyes. Lipid chain ordering is generally characterised by SCD parameter of lipids’ sn-1 and sn-2 methylene carbons. (85) The SCD order parameter can be expressed as (4), where *θ* is angle between normal to bilayer and vector joining the *Ci* to its hydrogen atom and < > represents ensemble and time average.

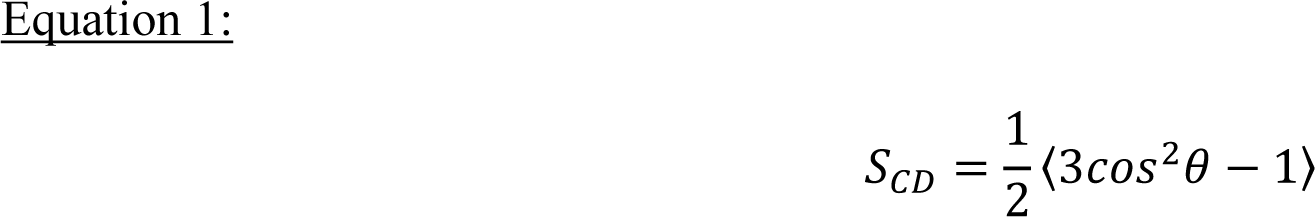

### 4AP-Cn dye staining

4AP-Cn (n=9,13) dye powders were dissolved in DMSO for working stock solutions of a concentration of 10mM. Working stocks are stable at room temperature, but were stored at 4°C, where they are stable for extended periods. Dyes were added to make the final concentration of 50μM in the bacterial culture solution and mixed by gentle pipetting. (Usage of prolonged vortexing for mixing causes a significant amount of the less integrated dye, 4AP-C9, to become unbound, so it must be avoided). After incubation at 37°C (without shaking) for 30 mins, the stained bacteria were washed with PBS to remove unbound dye. They were then centrifuged and resuspended in 0.5 ml 1xPBS for further steps. Quantification of the antibiotic effects was performed using the PMDR.

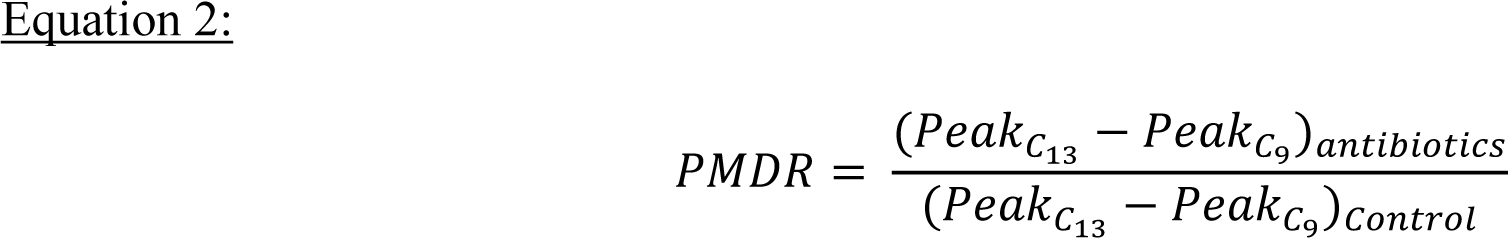

A meaningful PMDR will only arise if there exists a positive peak maxima difference in the control samples.

### Nile Red staining

Nile Red (Sigma Aldrich, St. Louis, Missouri, USA) in powder form was dissolved in DMSO to make a 1mg/ml stock, which was stored at −20°C. 2 µl of the stock solution per ml of bacterial solution in PBS was added, to make the final concentration in the solution to be 2 µg/ml. After keeping it at room temperature for 15 mins in the dark with no shaking, cells were washed with PBS to remove excess dye, centrifuged, and resuspended in 0.5 ml 1xPBS for further steps.

### CellRox Orange Staining

CellRox ® Orange (Invitrogen, Thermo Fisher Scientific), as a solution of concentration 2.5mM, was stored at −20°C. 2 µl of the dye was added per ml of the bacterial solution in PBS to make the final concentration in the solution to be 5 µM. According to the manufacturer’s directions, the cells were incubated for 30 mins at 37°C and washed with PBS to remove the excess dye. After that, the cells were centrifuged and resuspended in 0.5 ml 1xPBS for imaging and flow cytometry experiments.

Since the dye’s fluorescence upon oxidation is not very different from the exciting radiation, appropriate gating was performed to ensure that only positively stained cells are used to get the fold change of ROS induction. Positively stained cells are defined as cells whose median fluorescence index in the presence of the dye is greater than that corresponding to the top 0.1% of cells in the absence of the dye. This was further normalised by the fold change in untreated cells.

### DRAQ5 staining

DRAQ5 (BioLegend, San Diego, California, USA) as a solution of concentration 5mM was stored at 4°C for short- and long-term storage. 0.5 µl of the dye was added per ml of the bacterial solution in PBS to make the final concentration in the solution to be 2.5 µM. The cells were then kept in a shaking culture at 37°C for 30 mins, after which they were taken for confocal microscopy imaging.

### Laurdan Staining

Laurdan (Avanti Polar Lipids, Alabaster, Alabama, USA) in powdered form was dissolved in DMF to make a 10mM solution, with a working stock of 1mM in DMF, stored at −20°C. The dye was added (so that the final concentration of the dye in the solution is 10 µM) and mixed by vortexing for short periods. After incubation for 15 mins at 37°C, the stained bacteria were washed twice with 1x PBS supplemented with 0.2% Glucose and 1% DMF to remove the unbound dye, as in (86). They were resuspended in 0.5 ml supplemented PBS for further steps. Fluidity in quantitative terms is calculated using Laurdan Generalized Polarization (Laurdan GP). Laurdan GP is the intensity difference ratio at the two emission peaks, corresponding to its emission when surrounded by the fluid and gel-like phases (86). Here Laurdan GP is defined as follows (87) –

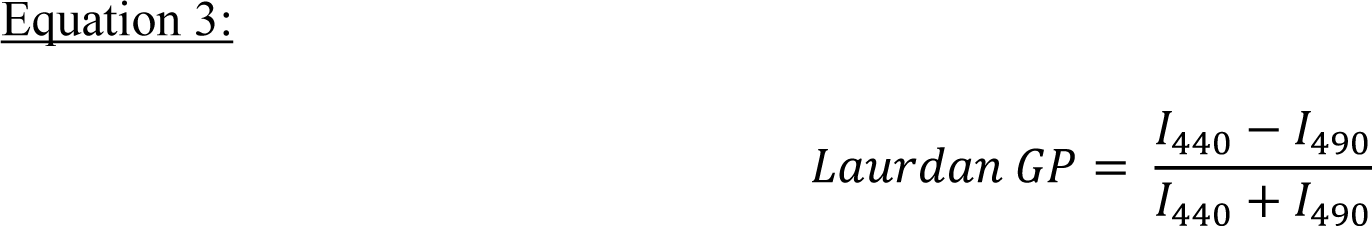

### BODIPY-C11 Staining

BODIPY™ 581/591 C11 (Invitrogen, Thermo Fisher Scientific) in powder form, was dissolved in DMSO, and then stored at −20°C. Since this dye behaves differently from the other dyes, staining was done prior to the secondary culture being made, allowing the stained cells to grow in the presence of the antibiotics and thereby pick up the lipid peroxidation when it occurs. After the primary culture was made, it was diluted to a solution of OD_600_ of 0.4, then incubated with a final concentration of 10μM of the dye at 37°C for 30 mins at 500rpm, before 100μl of this was used to inoculate the secondary culture containing the antibiotics. After growth, cells were pelleted down, washed with 1xPBS, then resuspended in 0.5 ml 1xPBS and then taken for flow cytometry measurements (38, 39, 88). The fluorescence of BODIPY-C11 upon activation is significantly distinct, so fluorescence in its active band can be assumed to be a direct result of peroxidation of the lipid membrane, which can then be normalised by the fold change of peroxidation in untreated cells.

### Fluorescence Spectroscopy Measurement

Fluorescence spectroscopy measurements were done primarily in two systems: sensitive experiments requiring high accuracies, like the dose-dependent antibiotic curves and the dye characterisation, were performed in a HORIBA JOBIN YVON Spectrofluorometer (FluoroMax – 4) (HORIBA Scientific, Edison, New Jersey, USA) in a fluorescence cuvette of path length 5 mm (Perkin Elmer B0631123), after putting 0.5ml of the sample solution in it. The excitation and emission slits (bandpass) were fixed at 5nm, with 1 nm increments in wavelength and an integration time of 0.1s. In this system, for the 4AP-Cn dyes, excitation was done at 375 nm, and the emission was observed from 400 nm to 700 nm. Most of the other experiments, like those with screening, clinical isolates, temperature-sensitive experiments, or those with Laurdan, were performed in a Tecan infinite 200 Pro plate reader (Tecan Group Ltd., Männedorf, Switzerland), with the samples in a flat bottom 96 well black plate. Experiments with Laurdan were performed similar to (86) with excitation at 350 nm, and an emission scan, with emission at 440 nm and 490 nm used for the GP calculation.

### Flow cytometry

Flow cytometry experiments were performed on the stained cells suspended in PBS. Measurements were performed in a BD FACS Canto® flow cytometer (BD Biosciences, Erembodegem, Belgium), with 50,000 events analysed per sample. Staining efficiency for Nile Red and 4AP-Cn dyes were compared, with 4AP-Cn dye fluorescence being measured in the V500 band (excitation – 405nm, emission – 485 to 535nm) and Nile Red in the PerCP-Cy5.5 band (excitation – 488nm, emission – 670LP). MFI fold changes were calculated for the quantification of ROS generation by CellROX Orange (Supplementary Figure 7) and lipid peroxidation by BODIPY-C11 (Supplementary Figure 8). CellROX Orange fluorescence was measured in the PE band (excitation – 488nm, emission – 564 to 606nm), while BODIPY-C11 was measured in the PE and FITC (excitation – 488nm, emission – 515 to 545nm) bands. Analysis was performed using the FlowJo software.

### Confocal Microscopy

After sample preparation, 20 to 30 µl of live cells were drop-cast on a frosted glass slide and covered with a cover slip before being sealed with transparent nail polish to prevent drying of the stained samples. Confocal Microscopy was done in a Zeiss LSM880 (Carl Zeiss AG, Oberkochen, Germany) in the bioimaging facility, at 100x oil immersion mode, with 3x optical zoom. Imaging in normal mode was performed to get the bright field images. Imaging for fluorescence visualisation was performed in AiryScan mode. Time series were collected for SRRF analysis.

### Super-resolution radial fluctuations (SRRF) analysis

Super-resolution radial fluctuations (SRRF) principle is similar to Single Molecule Localization Microscopy; the primary difference is that it does not depend upon the sensing of spatiotemporally isolated fluorescent molecules. However, SRRF introduces a magnified pixel grid called ‘subpixels’, and these ‘subpixels’ denote the probability of fluorescent molecules existing in that corresponding locality (89). To do this, SRRF measures the local radial symmetry, called radiality, in each pixel of the magnified image. A high radiality value corresponds to the presence of a fluorescent molecule at that position. Radiality fluctuations over time can be used to study the positions of the fluorescent molecule, but it is the degree of temporal correlations of the radiality that gives the final SRRF image. The SRRF algorithm was obtained as the NanoJ-SRRF plugin in ImageJ (90). SRRF analysis was performed by first splitting the channels when needed (corresponding to the two fluorophores) and then applying the SRRF analysis to the two independent channels. We used default parameters for SRRF analysis, given by 0.5, 5 and 6 ring radius, radiality magnification and axes in ring, respectively. Temporal radiality average (TRA) was used for temporal parameter. After the SRRF analysis, images are generated using ImageJ software using false colours for dye channels when needed.

### Image Analysis and Illustrations

Image analysis was performed using ImageJ, while illustrations were made using Inkscape and Microsoft PowerPoint.

## Data Analysis

Using Origin Pro 2018, noise from the fluorescence spectra data was removed by smoothening the curves using the Savitzky – Golay method (7-point window, 3 order polynomial). The smoothened data was then normalised to 0 to 1. Overall intensity changes are not relevant for the purposes of this study, and it is the relative differences in the spectra, that is, the wavelength at which the normalised intensity is equal to 1, which is referred to as the PMD, which is important for our purposes. Quantification via the generation of graphs and the subsequent significance tests were performed in GraphPad 8. The simultaneous comparison graph of ROS amounts and PMDRs across antibiotics in *E. coli* was generated using Plotly’s Python graphing library.

## Acknowledgements

We thank Dr H.C. Sudeeksha, Inorganic and Physical Chemistry Department, IISc, for assistance with fluorescence spectroscopy experiments, Prof. Jayanta Chatterjee from Molecular Biophysics Unit, IISc for providing *S. aureus* strains and members of the Bioimaging and Flow Cytometry facility at IISc for their technical help. This study was funded by a grant to D.N. on antibiotic resistance from the Department of Biotechnology, Government of India. The infrastructural support from the DBT-IISc program and DST-FIST grants are greatly acknowledged. We thank Prof. Rajendra Prasad and Prof. Amit Singh for facilitation this collaborative study and all previous and current members of the Nandi lab for their enthusiasm in supporting this investigation.

## Author Contributions

A.K.D. conceived the research, designed, and performed experiments and data analysis, interpreted results, and wrote the manuscript. T.V. conceived the research, aided with data interpretation, and helped with experiments. D.S. and B.K. performed MD simulations, experiments and data analysis. P.A. helped with data analysis. E.P.B. aided with clinical isolates and data analysis. S.S. and D.N. conceived the research and assisted in data interpretation and manuscript editing.

## Competing Interests

The authors declare no competing interests.

## Notes

### Competing Interest Statement

The authors have declared no competing interest.

