## Supplementary materials for "Cytoplasmic membrane thinning observed by interfacial dyes is likely a common effect of bactericidal antibiotics"

### Supplementary Materials – Figures and Tables

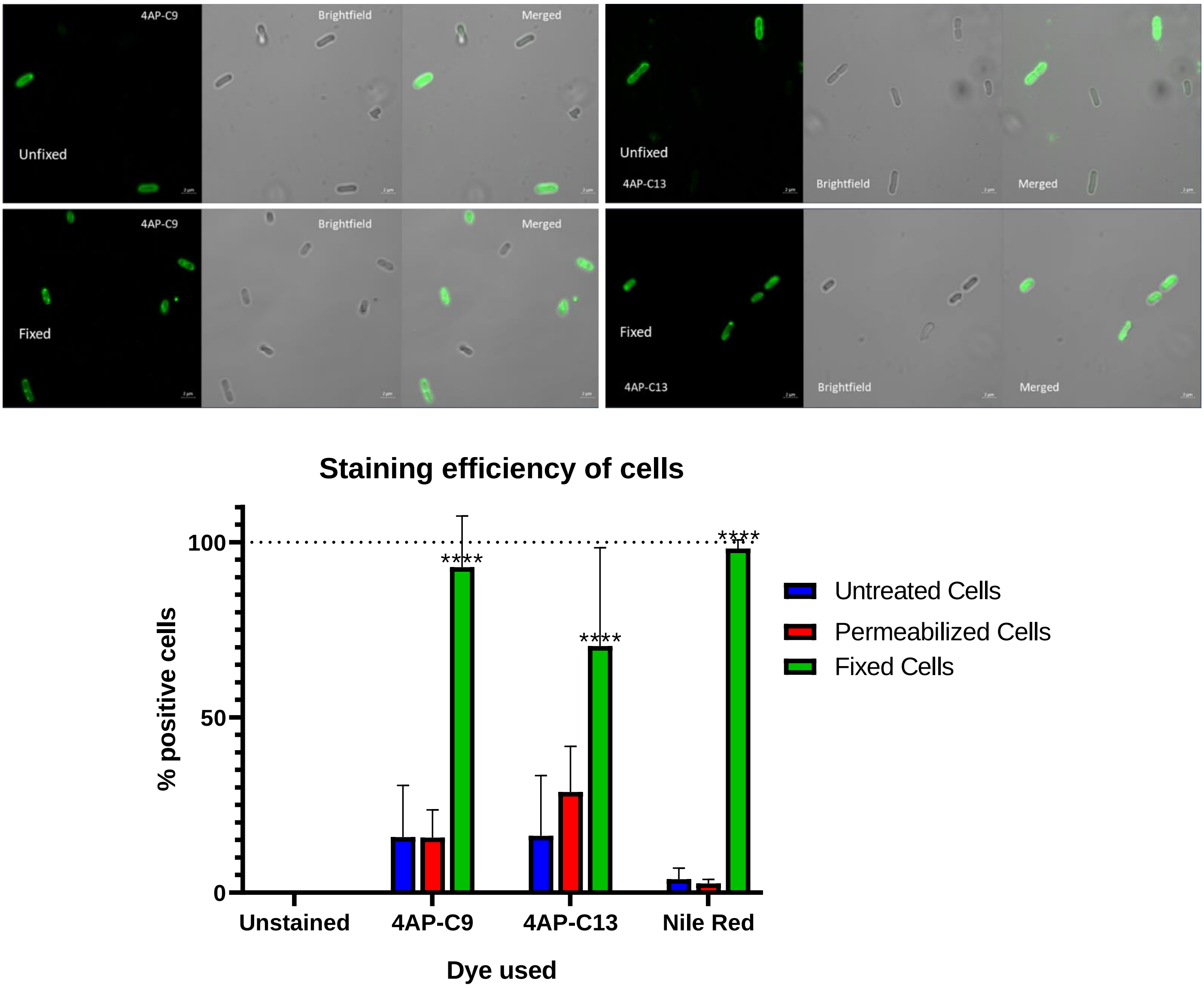


Supplementary Figure 1 – **Staining of Bacterial Cells is enhanced upon membrane permeabilization and cell functioning cessation**. Wild Type *E. coli* cells were grown for 6 hours and kept overnight at 4°C, after being subject to the different treatment conditions. For fixation, treatment with 4% PFA was performed before overnight storage, while permeabilization was performed via growth in the presence of 2µg/ml PMBN. While fixation was used as the determinant of permeabilization and cell functioning cessation, PMBN treatment was used as the determinant of outer membrane permeabilization. The staining efficiency of bacterial cells is visualized here for both dyes using confocal microscopy for untreated and fixed cells. Quantification of the staining efficiency in comparison to Nile Red has been performed using flow cytometry across untreated, permeabilized and fixed cells. The data shown here is representative of 10 biological replicates. Statistical analysis was performed here using two-way ANOVA, where * indicates $P < .05$; **, $P < .01$; ***, $P < .001$ and ****, $P < .0001$.

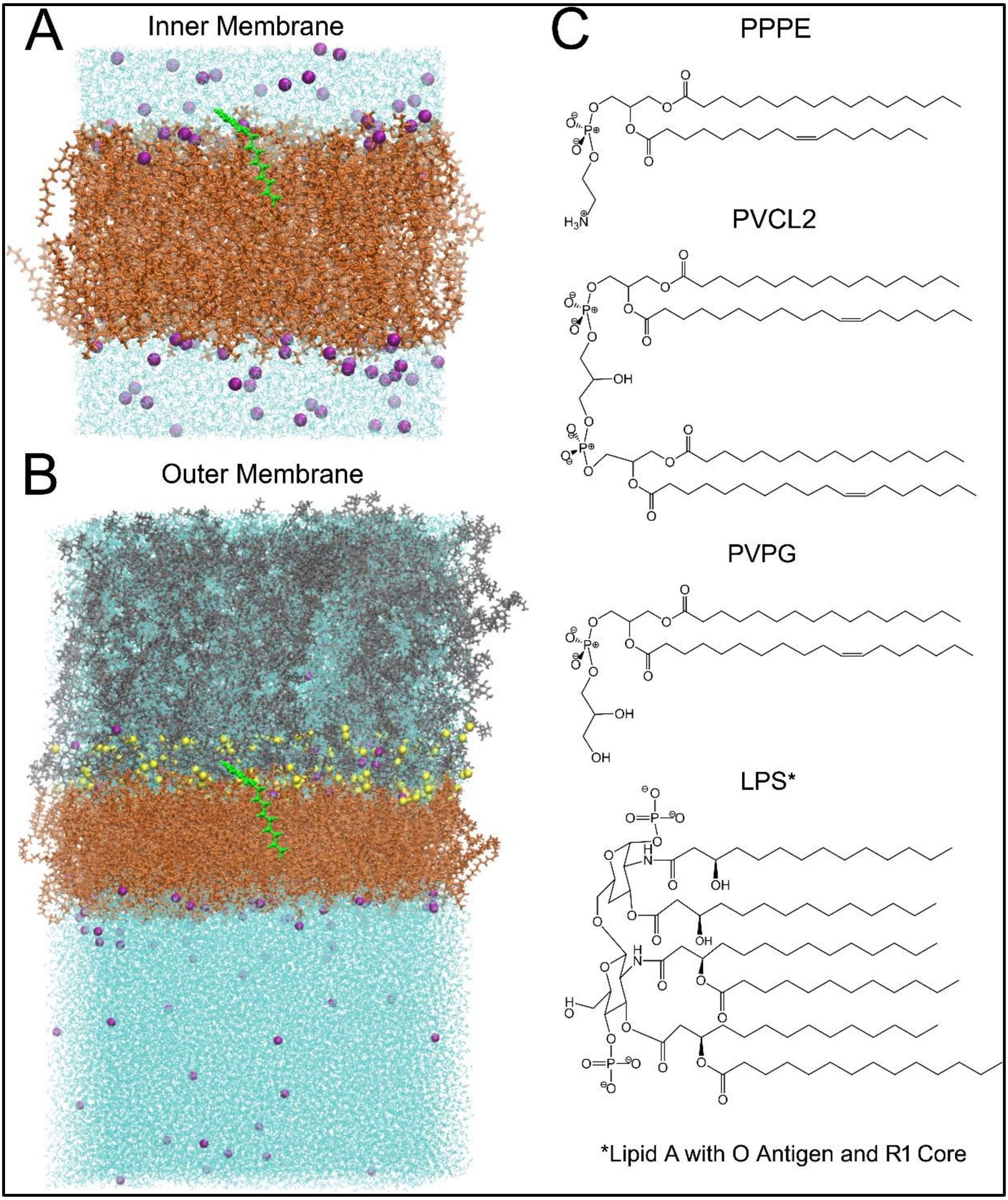


Supplementary Figure 2 – **Molecular dynamics simulations system snapshots and lipid compositions.** This figure shows the snapshots from MD simulations showing the most probable position of the 4AP-Cn dye (4AP-C9 shown for representation) inside the full simulated systems of (A) symmetric PL: PL and (B) asymmetric LPS: PL bilayer at 303.15 K. (C) The lipid structures used to create the bacterial IM and OM (details are given in Supplementary Table 1). Colour code used: Cyan – water molecules; purple – $K^+$ ions; yellow – Ca2+ ions; orange – alkyl tails of lipids; green – 4AP-Cn dyes; grey – LPS R1 core, which consists of (→3)bDGlc(1→3)[aDGal(1→2)aDGal(1→2)]aDGlc(1→3)aDGlc(1→3)[aLDHep(1→7)]aLDHep(1→3)aLDHep(1→5)[aDKdo(2→4)]

<table>
<tr><th colspan="5">Details of Molecular Dynamics simulations</th></tr>
<tr><td colspan="5">Composition of simulated lipid bilayers used in this study</td></tr>
<tr><th colspan="2" rowspan="2">Systems</th><th rowspan="2">Lipid Compositions</th><th colspan="2">Lipids</th></tr>
<tr><td>Upper Leaflet</td><td>Lower Leaflet</td></tr>
<tr><td colspan="2">LPS:PL [72:200]</td><td>LPS-PPPE/PVPG/PVCL2</td><td>72</td><td>150:40:10</td></tr>
<tr><td colspan="2">PL:PL [100:100]</td><td>PPPE/PVPG/PVCL2</td><td>75:20:5</td><td>75:20:5</td></tr>
<tr><td colspan="5">Details of simulated lipid bilayer/4AP-Cn systems</td></tr>
<tr><td>S. No</td><td>System</td><td>Temperature (K)</td><td>Total no. of atoms</td><td>Simulation length (ns)</td></tr>
<tr><td>1</td><td>IM (PL: PL)</td><td rowspan="6">303.15</td><td>51765</td><td>200</td></tr>
<tr><td>2</td><td>IM (PL:PL) + 4AP-C9</td><td>51810</td><td>500</td></tr>
<tr><td>3</td><td>IM (PL:PL) + 4AP-C13</td><td>51822</td><td>500</td></tr>
<tr><td>4</td><td>OM (LPS: PL)</td><td>302104</td><td>100</td></tr>
<tr><td>5</td><td>OM (LPS:PL) + 4AP-C9</td><td>302149</td><td>100</td></tr>
<tr><td>6</td><td>OM (LPS:PL) + 4AP-C13</td><td>302161</td><td>100</td></tr>
</table>

Supplementary Table 1 – **Details of the composition and conditions of the systems used for the molecular dynamics studies.** This table details the composition of the systems and the conditions under which the molecular dynamics studies

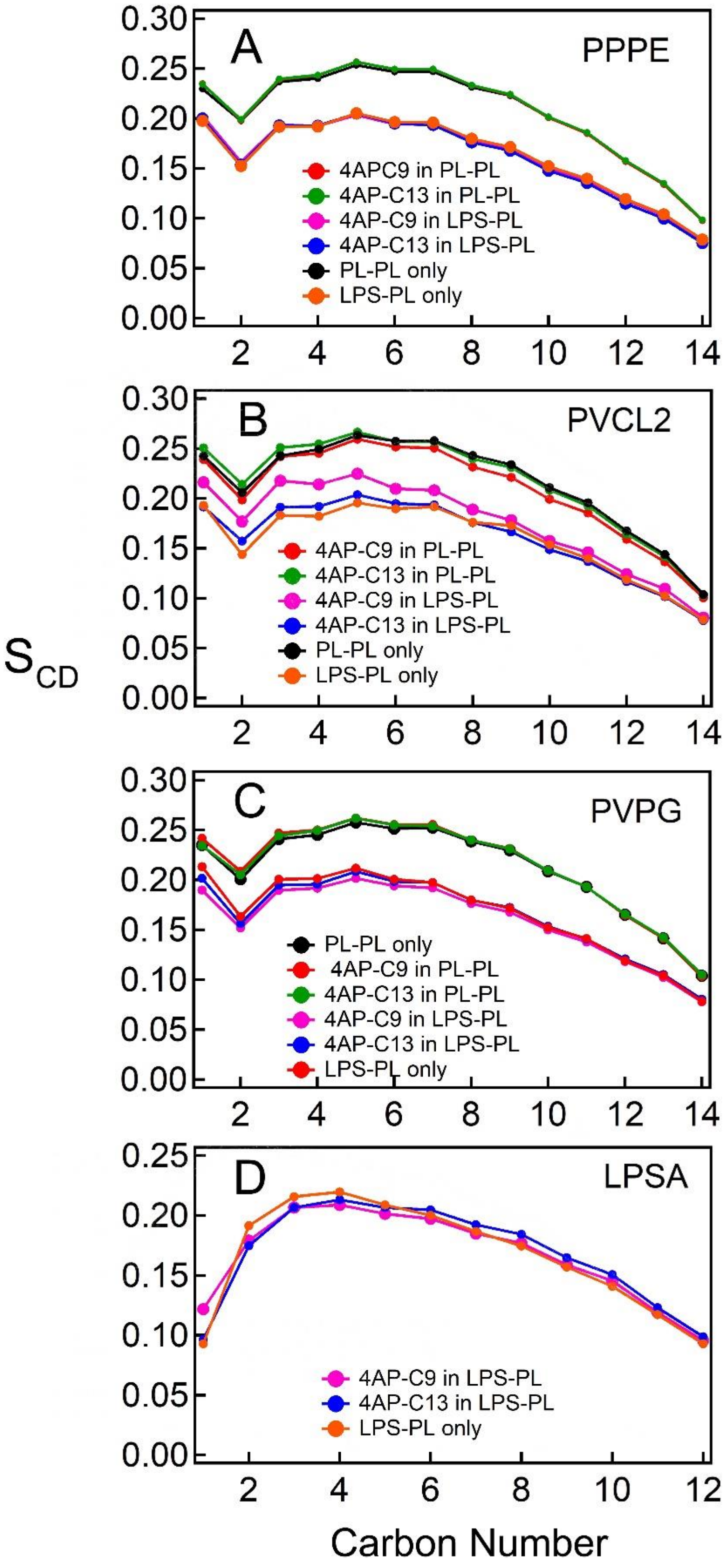


Supplementary Figure 3 – **Lipid chain order shows that the incorporation of 4AP-Cn dyes does not significantly affect the lipid order in IM or OM.** This figure shows the lipid chain order parameter ($S_{CD}$) obtained from MD simulations in the absence and presence of the 4AP-Cn dyes for (A) PPPE, (B) PVCL2, (C) PVPG and (D) LPSA. $S_{CD}$ for only one of the acyl chains of all the lipids has been shown here.

| **Antibiotics and their targets** | |
|---|---|
| Bactericidal antibiotics | |
| Polymyxin B | It consists of antimicrobial peptides (1, 2) which bind to the negatively charged LPS and replace the $Ca^{2+}$ and $Mg^{2+}$ cations, disrupting the outer membrane. (3, 4). Bacterial death likely occurs due to inner membrane disruption by PMB straddling it (4) or via binding to inner membrane LPS (5), among other reasons (6). PMB is known to induce ROS production (7-9). |
| Ciprofloxacin | It is a second-generation fluoroquinolone that primarily functions by inhibiting the bacterial type II topoisomerase (DNA gyrase) and the topoisomerase IV (10, 11), inhibiting cell division and blocking bacterial growth. (12) The bactericidal activity is dependent upon the generation of reactive products (ROS, RNI, etc.) (13-16) |
| Nitrofurantoin | It is an antibiotic that needs to be activated inside bacteria by reduction via the flavoprotein nitrofurantoin reductase to unstable metabolites, disrupting protein production and potentially damaging nucleic acids and other intracellular components. (17-20) |
| Ampicillin | It is a penicillin antibiotic, a type of β-lactam antibiotic. β-lactam antibiotics inhibit cell wall biosynthesis in bacteria and are the most used type of antibiotics (21-23). All β-Lactams share the highly reactive four-membered β-lactam ring and function by structurally mimicking the D-alanyl-D-alanine moiety of the peptidoglycan stem peptide to form an irreversible penicilloyl-β-lactam intermediate within the active site of the Penicillin Binding Proteins (PBPs) (24). Ampicillin, a pan-PBP inhibitor, irreversibly inhibits the transpeptidase enzyme and causes intracellular redox alterations (13, 25). |
| Amoxyclav (amoxicillin-clavulanic acid) | Amoxyclav is a combination of amoxicillin and clavulanic acid. Amoxicillin is a penicillin antibiotic similar to ampicillin, while clavulanic acid is a mechanism-based β-lactamase inhibitor, which reduces bacterial resistance to β-lactam antibiotics (21). |
| Ceftriaxone | It is a third-generation cephalosporin (26, 27), which is a separate class of structurally different β-lactam antibiotics, with a similar mode of action. |
| Meropenem | It is a carbapenem, which is another class of structurally distinct β-lactam antibiotics, albeit with a similar mode of action. Contrasting other β-lactams, they are highly degradation resistant to β-lactamase action (28), with a broader antimicrobial spectrum (29). |
| Gentamicin | It is an aminoglycoside, which are antibiotics that inhibit protein synthesis via various mechanisms, including binding with the bacterial membrane-associated ribosome, which impairs translational proofreading, leading to the formation of inaccurate proteins and eventually cell death. (30, 31) |
| Kanamycin | It is also an aminoglycoside and functions in a similar manner to gentamicin. (32) |
| Bacteriostatic antibiotics | |
| Tetracycline | It is an antibiotic that inhibits bacterial protein synthesis by blocking the attachment of charged aminoacyl-tRNA to the A site on the ribosome and by binding to the 30S and 50S subunits (33-35). |
| Azithromycin | It is a macrolide antibiotic that binds to the 50S subunit of the bacterial ribosome, thereby inhibiting mRNA translation and protein synthesis, among other uses. (36, 37) |
| Chloramphenicol | It is an antibiotic that inhibits protein synthesis by binding to the 50S ribosomal unit, preventing peptide bond formation directly, unlike macrolides which do so sterically. (38-40) |

Supplementary Table 2 – **Mechanisms of the different bactericidal and bacteriostatic antibiotics used.** This table details the known mechanisms of action of the various antibiotics used in this study, arranged by their bactericidal or bacteriostatic nature.

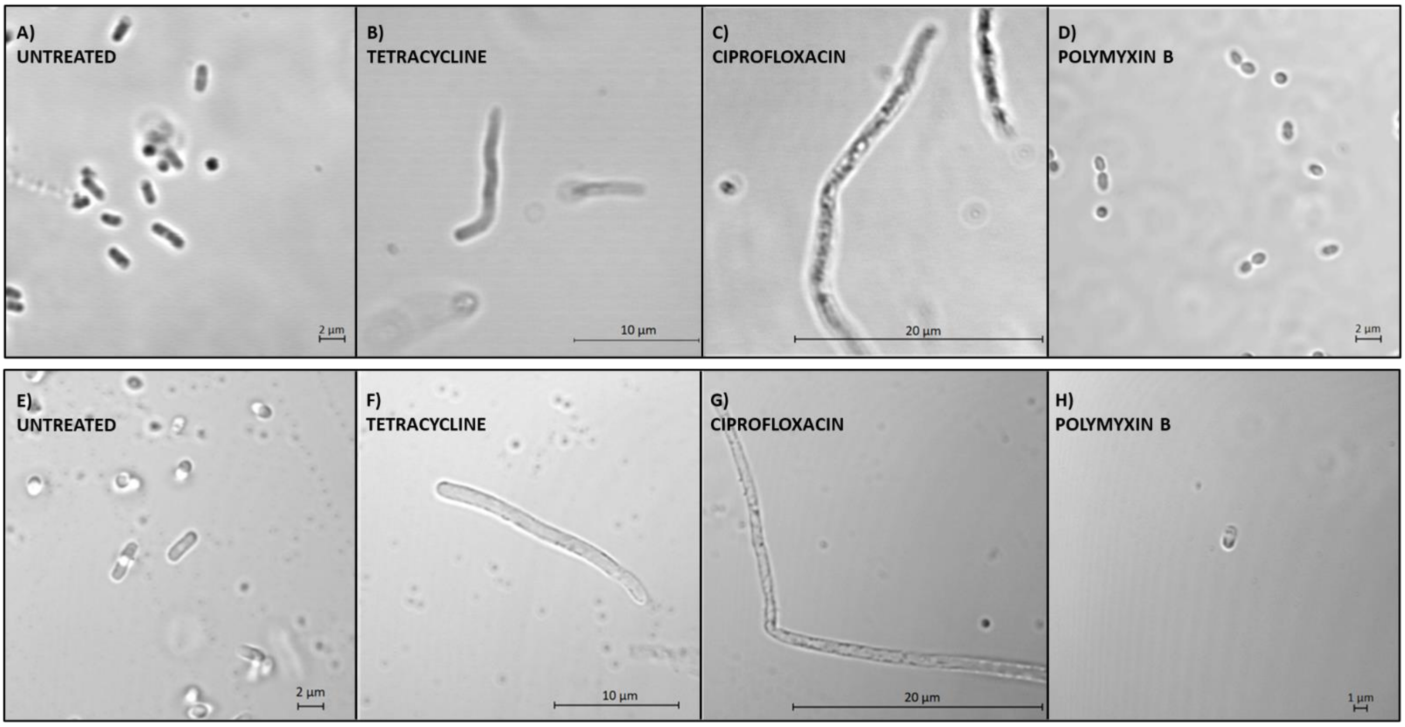


Supplementary Figure 4 – **Brightfield images of *E. coli* under different conditions of antibiotic treatment.** Brightfield images of the fields of view captured by confocal microscopy in Figure 4, are shown to allow comparison of the morphological changes as visualized in the presence and absence of dye staining. The images correspond to live Wild Type *E. coli* cells grown for 6 hours, which are (A, E) untreated or treated with 0.7XMIC of (B, F) Tetracycline, (C, G) Ciprofloxacin and (D, H) Polymyxin B.

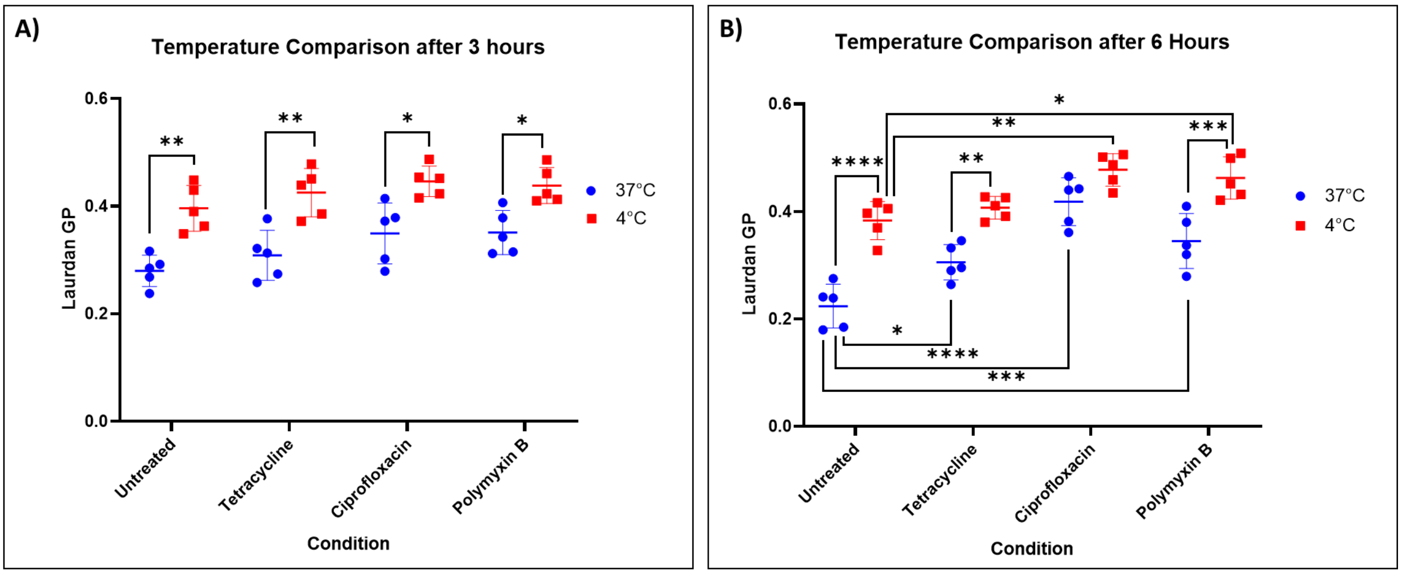


Supplementary Figure 5 – **Comparison of Fluidity across antibiotic treatment at low and high temperatures using Laurdan shows no measurable antibiotic induced increase in fluidity.** Laurdan GP is used to compare the fluidity changes in the membrane caused by the growth of the bacteria for (A) 3 hours and (B) 6 hours, in the presence of 0.7xMIC of the indicated antibiotic and compared to the untreated controls. The data shown here is representative of 5 biological replicates. Statistical analysis was performed here using two-way ANOVA, where * indicates $P < .05$; **, $P < .01$; ***, $P < .001$ and ****, $P < .0001$.

| Antibiotic Used | Concentration | Organism |
| --- | --- | --- |
| Azithromycin | 32 µg/ml | *E. coli* MG1655 |
| Tetracycline | 0.7 µg/ml | *E. coli* MG1655 |
| Chloramphenicol | 3 µg/ml | *E. coli* MG1655 |
| Polymyxin B | 180 ng/ml | *E. coli* MG1655 |
| Amoxicillin-Clavulanic Acid | 4 µg/ml | *E. coli* MG1655 |
| Ceftriaxone | 32 ng/ml | *E. coli* MG1655 |
| Ampicillin | 2 µg/ml | *E. coli* MG1655 |
| Ciprofloxacin | 24 ng/ml | *E. coli* MG1655 |
| Meropenem | 35 ng/ml | *E. coli* MG1655 |
| Nitrofurantoin | 6 µg/ml | *E. coli* MG1655 |
| Kanamycin | 10.5 µg/ml | *E. coli* MG1655 |
| Gentamicin | 5 µg/ml | *E. coli* MG1655 |
| Ciprofloxacin | 275 ng/ml | *Staphylococcus aureus* |
| Vancomycin | 3.2 µg/ml | *Staphylococcus aureus* |

Supplementary Table 3 – **Concentrations of the different antibiotics used, corresponding to ~0.7XMIC of mentioned bacterial strain.** The concentrations of the antibiotics to be used were determined by broth macro-dilution in our growth conditions to ensure an $OD_{600}$ ~0.2 on growth of the cells for 6 hours in the presence of the antibiotic.

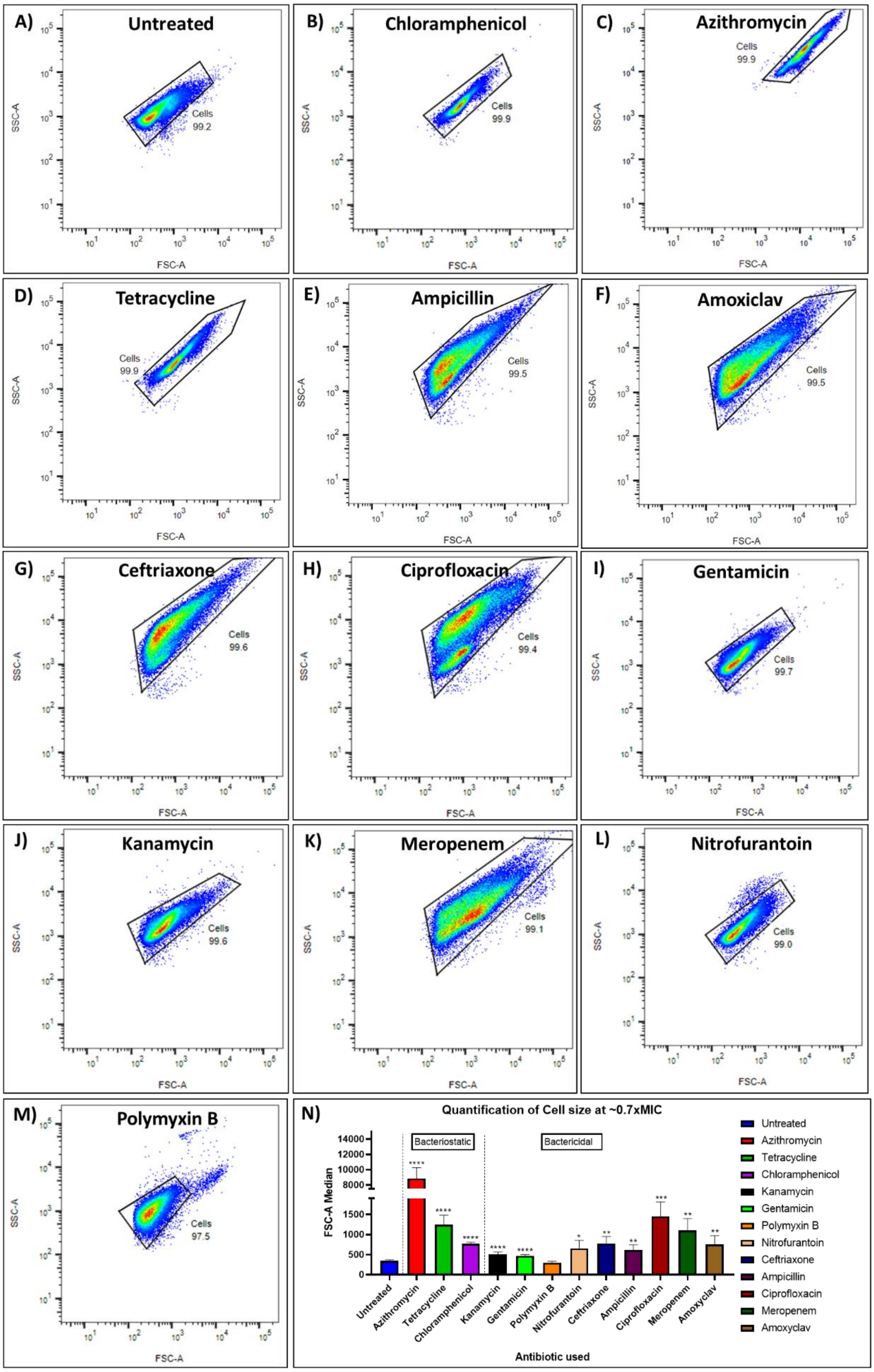
A) Untreated
Cells 99.2
B) Chloramphenicol
Cells 99.9
C) Azithromycin
Cells 99.9
D) Tetracycline
Cells 99.9
E) Ampicillin
Cells 99.5
F) Amoxiclav
Cells 99.5
G) Ceftriaxone
Cells 99.6
H) Ciprofloxacin
Cells 99.4
I) Gentamicin
Cells 99.7
J) Kanamycin
Cells 99.6
K) Meropenem
Cells 99.1
L) Nitrofurantoin
Cells 99.0
M) Polymyxin B
Cells 97.5
SSC-A
FSC-A
N) Quantification of Cell size at ~0.7xMIC
Bacteriostatic
Bactericidal
FSC-A Median
Antibiotic used
Untreated
Azithromycin
Tetracycline
Chloramphenicol
Kanamycin
Gentamicin
Polymyxin B
Nitrofurantoin
Ceftriaxone
Ampicillin
Ciprofloxacin
Meropenem
Amoxyclav

Supplementary Figure 6 – **The SSC-A vs FSC-A curve of *E. coli* is significantly different after treatment with antibiotics of different classes.** This figure shows the SSC-A vs FSC-A as obtained using flow cytometry, as a measure of the morphological changes for *E. coli* Wild Type MG1655 cells grown for 6 hours in the presence of ~0.7xMIC of antibiotics – A) untreated cells, B) Chloramphenicol, C) Azithromycin, D) Tetracycline, E) Ampicillin, F) Amoxiclav, G) Ceftriaxone, H) Ciprofloxacin, I) Gentamicin, J) Kanamycin, K) Meropenem, L) Nitrofurantoin and M) Polymyxin B. N) This figure compares the Median FSC-A for all the different antibiotic treatments, as a possible measure of cell size change, after treatment with the antibiotics. FSC-A represents forward scattering, while SSC-A represents side scattering. Data in (N) is representative of 8 replicates. Statistical analysis was performed here using Brown-Forsyth and Welch ANOVA test, where * indicates $P < .05$; **, $P < .01$; ***, $P < .001$ and ****, $P < .0001$.

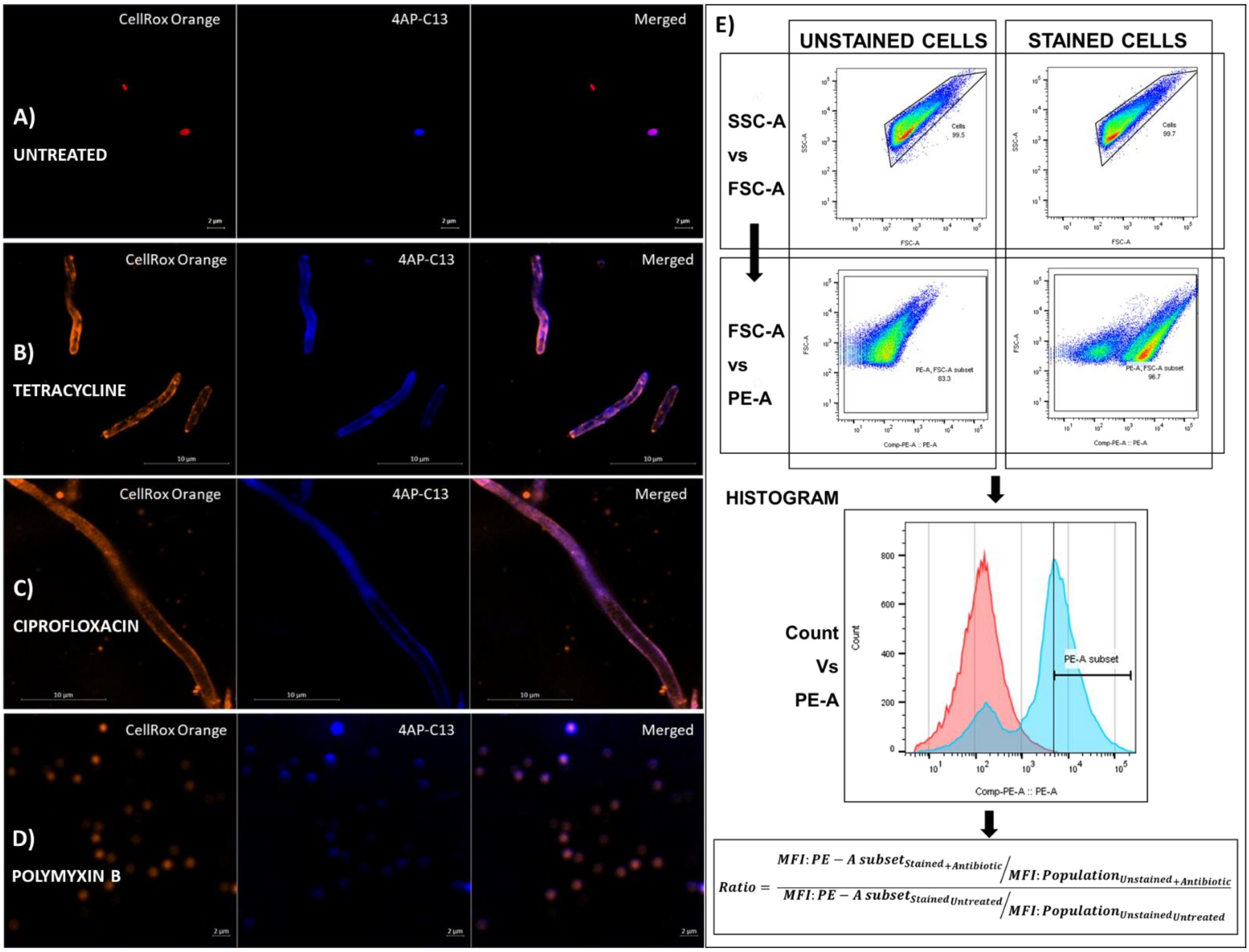


Supplementary Figure 7 – **Reactive oxygen species tend to congregate near the cell membrane, and their amounts on antibiotic treatment can be quantified using flow cytometry**. Confocal Microscopy (in AiryScan mode) was used to visualize the reactive oxygen species localization using CellROX Orange and 4AP-C13 in cells under differing treatments – A) untreated cells, and 0.7XMIC of B) Tetracycline, C) Ciprofloxacin and D) Polymyxin B. In (A), (B), (C) and (D), column 1 and column 2 show the dyes as separate channels, while column 3 shows both channels together, where the colocalization of the two dyes is made abundantly clear. E) This figure describes the methodology used for the quantification of ROS amounts for the different antibiotic-treated cells, via the usage of the Amoxyclav (amoxicillin-clavulanic acid) treated cells as an example.

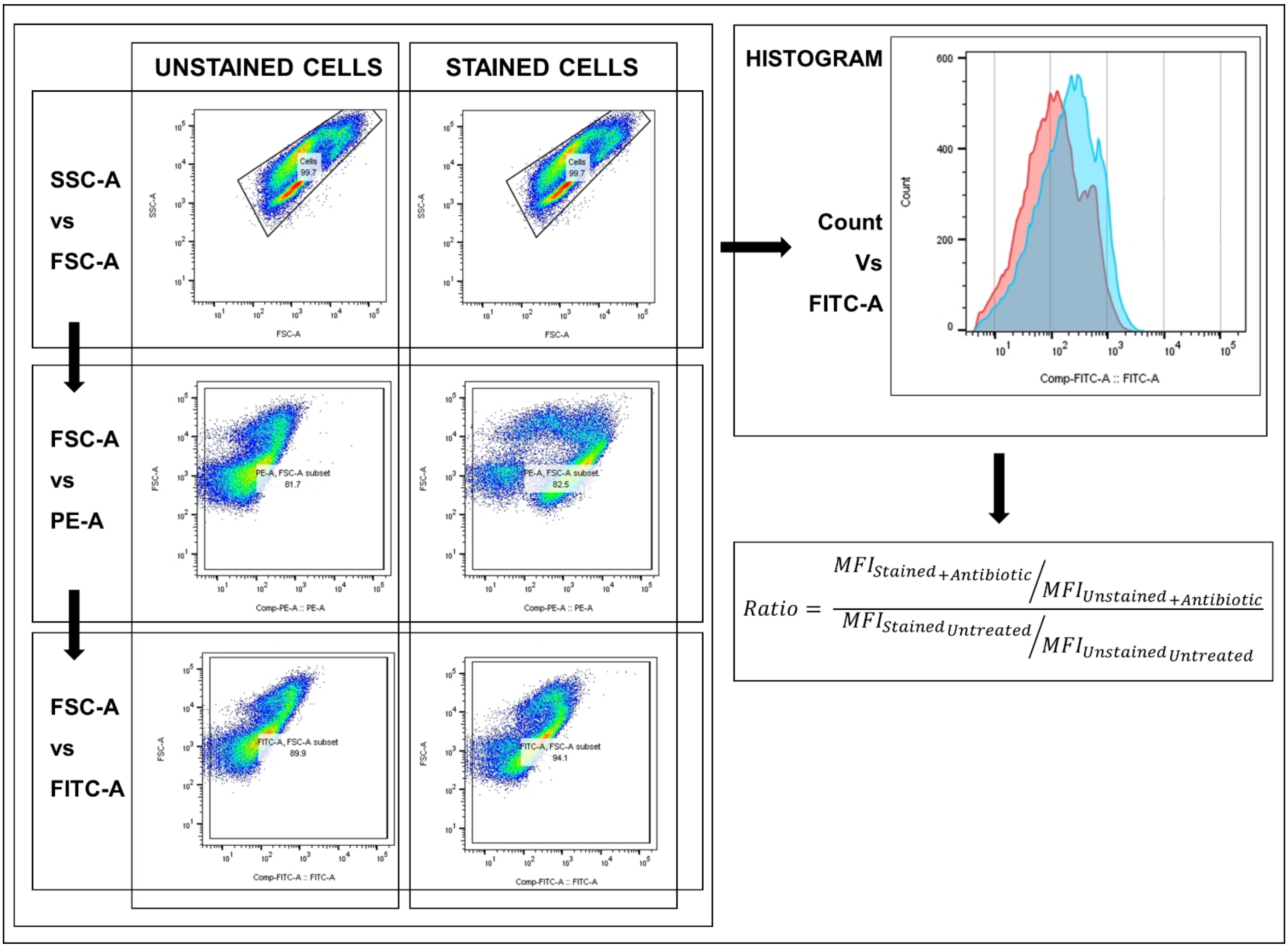


Supplementary Figure 8 – **Methodology for the quantification of lipid peroxidation on antibiotic treatment.** This figure shows the step-by-step methodology followed for the quantification and comparison of the extent of lipid peroxidation on different antibiotic treatments, using the ciprofloxacin treated cells as an example.

| Clinical Isolates | | | | |
|---|---|---|---|---|
| **Antibiotic** | **The concentration of antibiotics used (µg/ml)** | | | |
| | ***E. coli*** | ***K. pneumoniae*** | ***Enterobacter spp.*** | ***A. baumannii*** |
| Ciprofloxacin | 0.5 & 8 | 64 | 2 | 0.5 & 8 |
| Meropenem | 2 | 4 | 2 | 2 |
| Amoxyclav | 8 | 64 | - | - |
| Nitrofurantoin | 64 | 64 | 32 | - |
| Gentamicin | 16 | 4 | 16 | 2 & 8 |
| Polymyxin B | 0.5 | - | - | 0.5 |

Supplementary Table 4 – **Concentrations of the different antibiotics used during the growth of the clinical isolates of the indicated bacterial species.** The concentrations of the antibiotics to be used and the time of growth for a bacterial species were determined by broth macro-dilution in 96 well plates, with the sample preparation being performed on the growth of the cells in the presence of the antibiotic for 6 hours in the case of *E. coli* and *Enterobacter spp.* and 8 hours for *K. pneumoniae* and *A. baumannii* in 50 ml cultures in 150ml flasks.

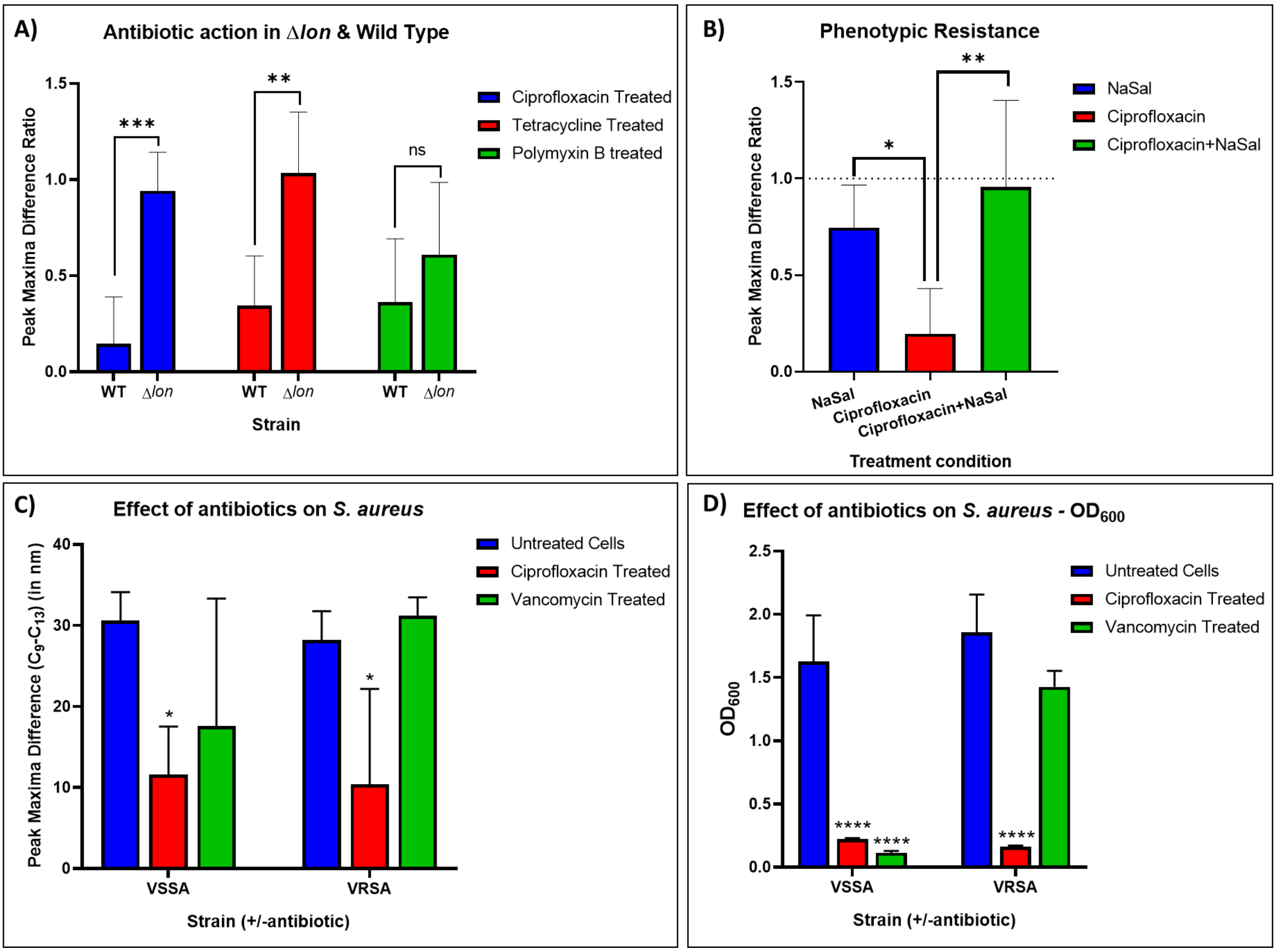


Supplementary Figure 9 – **Peak maxima difference ratio (PMDR) reduction is specific to sensitive bacteria, across bacterial species and modes of resistance.** (A) This figure shows the PMDR of the *E. coli* mutant strain Δ*lon* grown for 6 hours in the presence of the indicated antibiotics when compared with the Wild Type *E. coli* MG1655 strain under similar treatment conditions, as a measure of genotypic resistance. (B) This figure shows the effect of the phenotypic antibiotic resistance inducer sodium salicylate (NaSal) on ciprofloxacin treatment of Wild Type *E. coli* as seen by the rescue of the PMDR. Vancomycin Sensitive *Staphylococcus aureus* (VSSA) and Vancomycin-Resistant *Staphylococcus aureus* (VRSA) were grown for 6 hours in the absence and presence of ciprofloxacin and vancomycin. The peak maxima difference has been visualized in (C) while the $OD_{600}$ of the strains has been shown in (D). Data sets in (A) and (B) are representative of 7 replicates, while those in (C) and (D) are representative of 5 replicates. Statistical analysis was performed for (A), (C) and (D) using two-way ANOVA and (B) using one-way ANOVA, where * indicates $P < .05$; **, $P < .01$; ***, $P < .001$ and ****, $P < .0001$.

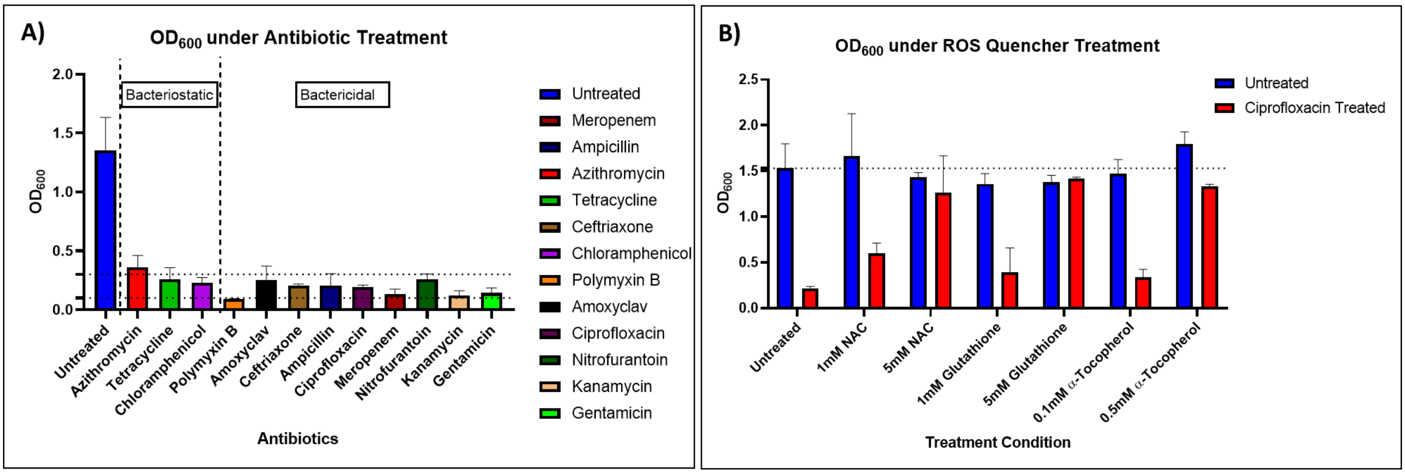


Supplementary Figure 10 – **$OD_{600}$ of relevant experimental conditions in lab strain *E. coli* MG1655.** This figure shows the $OD_{600}$ of the lab *E. coli* MG1655 under treatment of A) 0.7XMIC of various antibiotics and B) ROS quenchers in the presence and absence of Ciprofloxacin. The data in (A) and (B) are representative of 7 and 5 replicates, respectively.

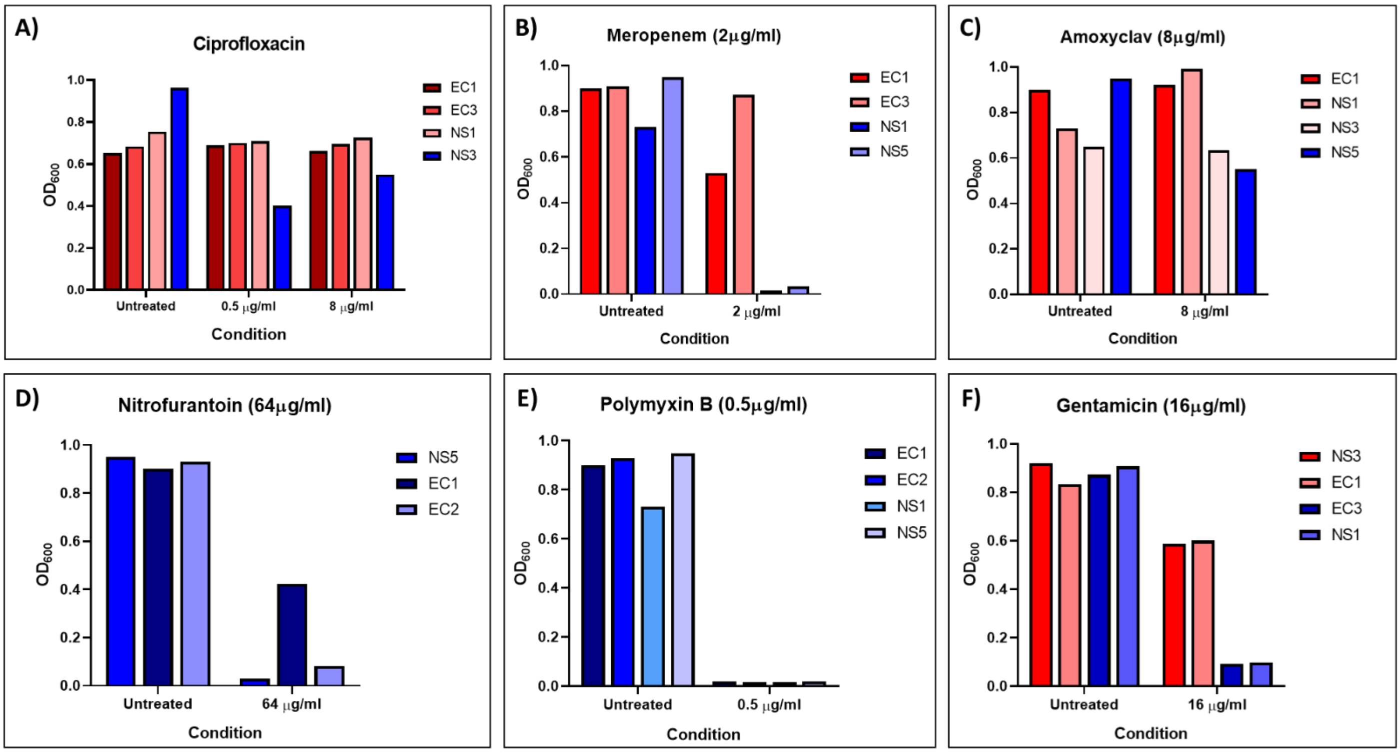


Supplementary Figure 11 – **$OD_{600}$ of *E. coli* clinical isolates under different antibiotic treatments**. The $OD_{600}$ of the various *E. coli* clinical isolates used in the study have been visualized, in the presence of different concentrations of the antibiotic – A) Ciprofloxacin (0.5 and 8µg/ml), B) Meropenem (2µg/ml), C) Amoxyclav (8µg/ml), D) Nitrofurantoin (64µg/ml), E) Polymyxin B (0.5µg/ml) and F) Gentamicin (16µg/ml). The clinical isolates marked red are resistant to the antibiotic, while those marked blue are sensitive. Sensitivities and resistances are designated based on $OD_{600}$ reductions of the isolate on antibiotic treatment – >~50% reduction in the presence of the antibiotic compared to its absence is designated sensitive.

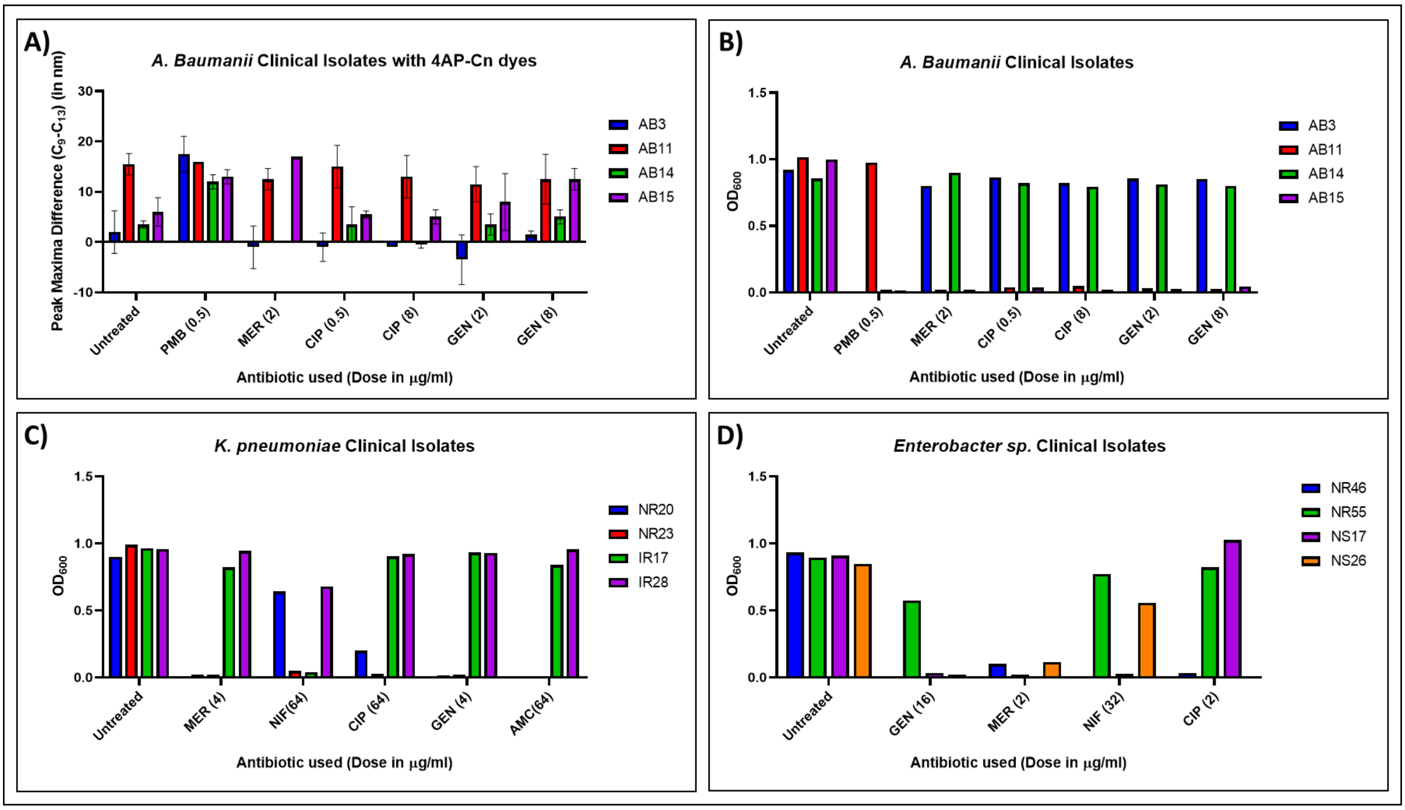


Supplementary Figure 12 – **Peak maxima difference (PMD) of *A. baumannii* isolates, and the $OD_{600}$ of *A. baumannii, K. pneomoniae* and *Enterobacter spp.* under different antibiotic treatments**. (A) This figure shows the PMDs for the clinical isolates of *A. baumannii*, while the $OD_{600}$ of the isolates under the different treatments is shown in (B). The $OD_{600}$ of the different *K. pneomoniae* clinical isolates in the presence of different concentrations of the antibiotic used in the study have been visualized in (C), while those of the *Enterobacter spp.* samples have been visualized in (D). For the PMDs visualized in (A), each data point is representative of 2 technical replicates.

---
